# Global, multiplexed dendritic computations under *in vivo*-like conditions

**DOI:** 10.1101/235259

**Authors:** Balázs B Ujfalussy, Máté Lengyel, Tiago Branco

## Abstract

Dendrites integrate inputs in highly non-linear ways, but it is unclear how these non-linearities contribute to the overall input-output transformation of single neurons. Here, we developed statistically principled methods using a hierarchical cascade of linear-nonlinear subunits (hLN) to model the dynamically evolving somatic response of neurons receiving complex spatio-temporal synaptic input patterns. We used the hLN to predict the membrane potential of a detailed biophysical model of a L2/3 pyramidal cell receiving *in vivo*-like synaptic input and reproducing *in vivo* dendritic recordings. We found that more than 90% of the somatic response could be captured by linear integration followed a single global non-linearity. Multiplexing inputs into parallel processing channels could improve prediction accuracy by as much as additional layers of local non-linearities. These results provide a data-driven characterisation of a key building block of cortical circuit computations: dendritic integration and the input-output transformation of single neurons during *in vivo*-like conditions.

## Introduction

Cortical neurons receive and integrate thousands of synaptic inputs within their dendritic tree to produce action potential output. A large repertoire of biophysical mechanisms supporting a remarkable diversity of input integration properties has been described in dendrites, including synaptic saturation (Abrahamsson et al., 2012), dendritic spikes (Häusser et al., 2000), NMDA receptor non-linearities (Major et al., 2013), and interactions between excitation and inhibition (Palmer et al., 2012). A fundamental function of these mechanisms is to control the way single neurons map spatio-temporal patterns of inputs, the spike trains impinging on their dendritic trees, to somatic membrane potential responses, and ultimately, action potential output. This input-output mapping at the level of individual neurons has a critical role in shaping the population dynamics and computations that emerge at the level of neural circuits (Rubin et al., 2015; Zador, 2000), and has been the focus of intensive investigation (Gerstner & Naud, 2009; Silver, 2010). Yet, the way dendritic inputs are mapped into somatic output under *in vivo* conditions remains poorly understood.

Despite considerable progress in characterising non-linear dendritic mechanisms, our understanding of neuronal input integration is limited, as it is mostly based on data from *in vitro* experiments studying neurons under simplified input conditions. These studies have characterised the fundamental properties of dendritic integration (e.g. the threshold and propagation of dendritic spikes) by parametrically varying a small set of input features, such as the number, location, and timing of inputs in periodic trains of synaptic stimulation (Losonczy & Magee, 2006; Branco et al., 2010; Branco & Häusser, 2011; Makara & Magee, 2013). Similarly, most theoretical approaches have focused on simplified input regimes, in which single neurons receive a small number of inputs (Hao et al., 2009; Jadi et al., 2012; Behabadi et al., 2012; Gidon & Segev, 2012), or their inputs and output have stationary firing rates (Poirazi et al., 2003b). However, dendritic integration *in vivo* can in principle exhibit different properties than *in vitro* because of the high density and complexity of the synaptic input patterns characteristic of *in vivo* states, and the high conductance regime they generate (London & Segev, 2001; Destexhe et al., 2003).

Recent experimental work has demonstrated that dendritic non-linearities can substantially contribute to neuronal output *in vivo* (Xu et al., 2012; Lavzin et al., 2012; Palmer et al., 2014; Takahashi et al., 2016), but it is still unclear whether active dendrites qualitatively change the neuronal input-output transformation. For example, while theoretical and experimental studies have suggested that voltage-gated non-linearities change the single neuron input-output fuction from linear to supralinear (Poirazi et al., 2003b; Polsky et al., 2004; Branco & Häusser, 2011; Makara & Magee, 2013), it is also possible that their role is more consistent with a re-weighting of inputs (Magee, 2000; Häusser, 2001), such that the relative contributions of different synapses are modified, while leaving the global computation performed by the individual neuron relatively unchanged (Cash & Yuste, 1998). This gap in our knowledge is in part due to shortcomings of currently available techniques used to characterise the inputoutput transformation of neurons in the low-input regime, which do not scale up to the *in vivo* situation where the neuron is constantly bombarded with thousands of presynaptic spikes. Thus, understanding the contribution of dendritic mechanisms for single neuron computations requires both technical advances that allow experimental measurements of the spatio-temporal dynamics of synaptic activation across entire dendritic trees *in vivo* (Jia et al., 2010; Wilson et al., 2016), and new analysis methods for describing and quantifying dendritic and single neuron computations.

To develop a new framework for analyzing single neuron input-output transformations, we take inspiration from the domain of sensory processing, where statistical models have been successfully applied to predict neuronal responses to sensory stimuli with complex spatio-temporal structure *in vivo* (Pillow et al., 2008; Ramirez et al., 2014). In these studies, the transformation of external inputs (e.g. visual images) to the neuronal response (e.g. of a visual cortical neuron) is expressed as a linear filtering step followed by a non-linear transformation (linear-nonlinear, LN models). This framework has the advantage that it allows the application of principled statistical methods to fit models directly to *in vivo* recordings, and yields easily interpretable functional descriptions - features that are both missing from approaches that aim to fit complex multicompartmental models to the data (Druckmann et al., 2007; Keren et al., 2009). However, in its standard form, the LN framework starts from sensory stimuli as the main input to the model, and so the recovered non-linearities are a combination of the non-linear processing steps of the upstream circuitry and the non-linearities implemented by the particular cell whose output is recorded and fitted (Antolik et al., 2016). Thus, for dissecting the contribution of single neuron processing, the LN framework needs to be endowed with a unique combination of features: inputs must solely be the spike trains received directly by the cell (Truccolo et al., 2010), the output must be its somatic response (Mensi et al., 2012; Ramirez et al., 2014), and a cascade of non-linear input-output transformations must be allowed (Vintch et al., 2015; Freeman et al., 2015) to account for various forms of non-linear processing in dendrites and the soma.

Here we combine these features and present a hierarchical LN model (hLN) that predicts the subthreshold somatic response of neurons to complex spatio-temporal patterns of synaptic inputs. We use hLN models to study dendritic integration in biophysically detailed compartmental models of a L2/3 cortical pyramidal cell and a cerebellar granule cell, that reproduce the main features of dendritic and somatic voltage activity recorded *in vivo* (Smith et al., 2013; Duguid et al., 2012). Surprisingly, we find that more than 90% of the somatic response can be accurately described by linear integration followed a single global non-linearity, and that capturing *in vivo* membrane potential dynamics can require a novel form of input processing, whereby dendritic subunits multiplex inputs into parallel processing channels with different time constants. For both pyramidal and granule cells we recover slow and fast processing channels with biophysical equivalents, such as synaptic integration via NMDA and AMPA receptors. Our approach provides a quantitatively validated and intuitive description of dendritic information processing performed by neurons receiving large barrages of synaptic inputs, and thus paves the way for understanding the role of dendrites in the computations performed by neuronal circuits.

## Results

### Responses to simple stimuli do not predict responses to complex stimulation patterns

In order to illustrate the shortcomings of the most widely used approach for characterising dendritic non-linearities (Polsky et al., 2004; Losonczy & Magee, 2006; Branco et al., 2010; Abrahamsson et al., 2012; Makara & Magee, 2013), we first implemented a previously validated multicompartmental biophysical model of a L2/3 cortical pyramidal cell (Smith et al., 2013) stimulated with inputs that were either similar to those typically used in *in vitro* experiments, or resembled naturalistic patterns expected to emerge *in vivo* (Experimental Procedures). We first probed the dendrite with regular synaptic stimulation trains of multiple (1-40) stimuli (1 stimulus per synapse, with 1 ms delays between synapses) while recording the somatic membrane potential response (Figure 1A). We then characterized the input-output non-linearity of the cell by comparing the magnitude of the measured response with that expected from linearly summing the responses to the individual synaptic stimuli (Figure 1B). To demonstrate the effects of non-linear dendritic processes in shaping this input-output non-linearity, we changed the NMDA-to-AMPA channel ratio (NAR) in the model. In line with previous results (Behabadi et al., 2012), higher NAR resulted in a marked increase in the supralinearity of the input-output transformation as revealed by this analysis method (compare red, purple and blue). However, the same model neurons showed very little difference in their behavior when stimulated with *in vivo*-like naturalistic input patterns (Figure 1C): apart from a slight tonic offset, their somatic response was very highly correlated (Figure 1D, top, R^2^=0.94).

**Figure 1.**
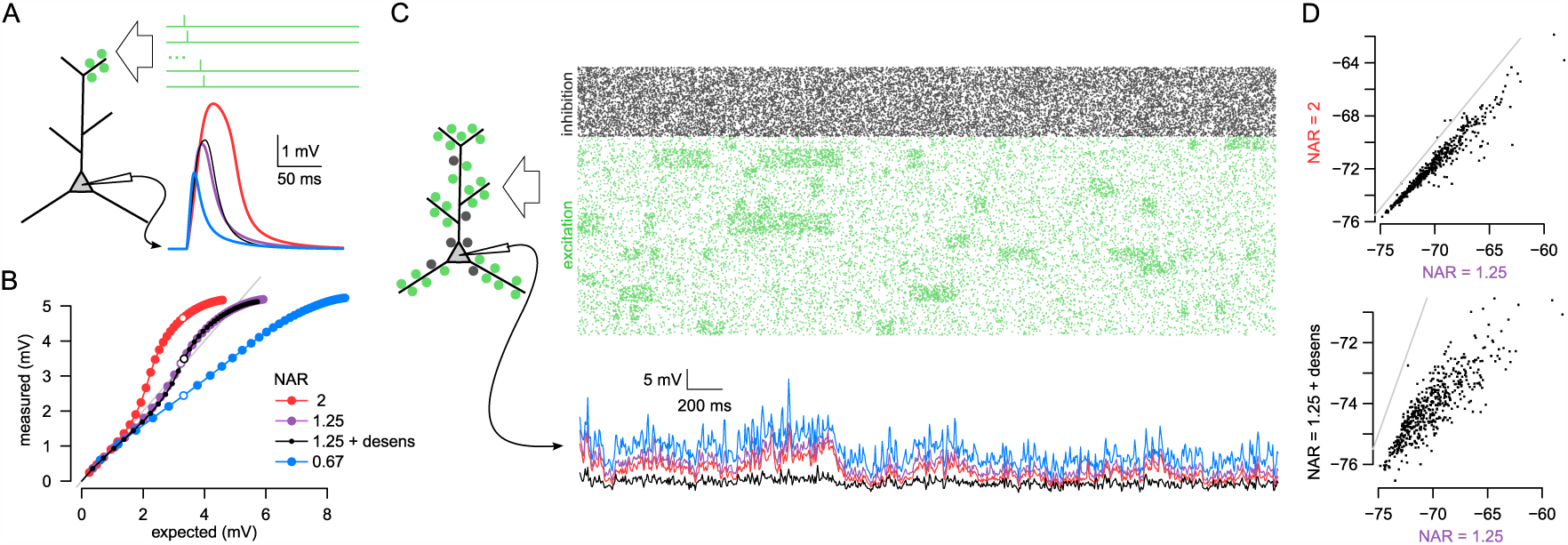
*In vitro* measurements of dendritic integration do not predict the response to *in vivo*-like synaptic activation patterns. (A) Illustration of a typical *in vitro* protocol stimulating a small number of neighbouring synapses (left) using a fixed temporal delay (top right) while recording the somatic membrane potential response (bottom right). Somatic responses of four model neurons are shown (colors as in B-C) to stimulus-sequences with the number of stimuli chosen such that the expected responses of all neurons were identical (see also circles in B). (B) Measured response amplitudes in different model neurons (colors) to the stimulation of 1-40 neighbouring synapses at 1 ms intervals as a function of the amplitude of the expected response. Open symbols indicate simulations shown in A (expected amplitude ~ 3.3 mV), gray line shows identity line (exactly linear integration). Models differed in the NMDA-to-AMPA ratio (NAR) of their glutamatergic synapses, resulting in qualitatively different modes of dendritic integration: superlinear (red, NAR=2), approximately linear (purple and black, NAR= 1.25) and sublinear (blue, NAR=0.67). Only the model shown in black included desensitization of NMDA receptors. (C) Responses of the same four model neurons as shown in A-B to sustained *in vivo*-like inputs with complex spatio-temporal structure. Top: input spike trains arriving at excitatory (green) and inhibitory synapses (gray). Bottom: the somatic membrane potential in the four neurons in response to the stimulus shown above (color code as in A-B). (D) Correlation between the responses of selected pairs of model neurons shown in C: neuron with NAR=2 (red) vs. neuron with NAR=1.25 (purple, top) and the neuron with desensitizing NMDA receptor (desens, black) vs. the neuron without NMDA desensitization (purple, both with NAR=1.25, bottom). Gray lines show identity lines. Note lack of relationship between correlations under *in vivo*-like conditions (D) and similarity of responses to in *vitro* stimulation protocols (A-B).

Conversely, while the simple *in vitro*-like stimulation protocol revealed no difference between two model neurons that differed in the kinetics of NMDA receptor desensitization (Figure 1A-B, purple vs. black), the same neurons showed markedly different responses to sustained inputs (Figure 1C) that were only weakly correlated (Figure 1D, bottom, R^2^=0.69). This was because the simple *in vitro* protocol only stimulated each synapse once, and therefore post-activation desensitization had no effect on the response, while *in vivo*-like sustained stimulation led to a difference in the fraction of receptors in the desensitized state between the two neurons.

These examples demonstrate that due to the dynamic and sustained nature of synaptic activation, differences and similarities across different cells under *in vivo*-like conditions cannot be readily predicted from their responses to simple *in vitro* stimulation protocols (London & Segev, 2001). Moreover, even under sustained stimulation, the degree to which dendritic processing appears nonlinear can depend on whether the stimulus is stationary, using constant firing rates (Poirazi et al., 2003b), or it shows fluctuations probing the full dynamic range of the neuron (Figure S1). This motivated us to study neuronal input-output transformations directly under *in vivo*-like conditions with complex spatio-temporal input patterns, rather than trying to extrapolate from their responses to simplistic stimuli. As the measured-vs-expected method does not generalise to this input regime, we developed a new model-based analysis technique to address this problem.

### Fitting the input of a biophysical model to *in vivo* dendritic recordings

Studying the contribution of dendritic non-linearities to the input-output mapping performed by neurons under realistic, *in vivo*-like conditions requires, ideally, observing both the inputs and the output of a particular neuron simultaneously, with high spatio-temporal precision. Although the combination of high resolution two-photon imaging techniques with *in vivo* patch clamp recordings will likely deliver this sort of data in the near future (Grienberger & Konnerth, 2012), such data is not yet available. Therefore we took a two-step approach (Figure 2A): first, we used a detailed biophysical model neuron to reproduce dendritic activity recorded experimentally *in vivo* (Figure 2A, fit 1), and then used the known inputs and outputs of the biophysical model to study the input-output transformation of single neurons under *in vivo*-like input conditions (Figure 2A, fit 2). Crucially, we paid careful attention to match our biophysical model as closely as possible to *in vivo* data to ensure that the non-linearities measured *in vivo* were also expressed by our model.

**Figure 2.**
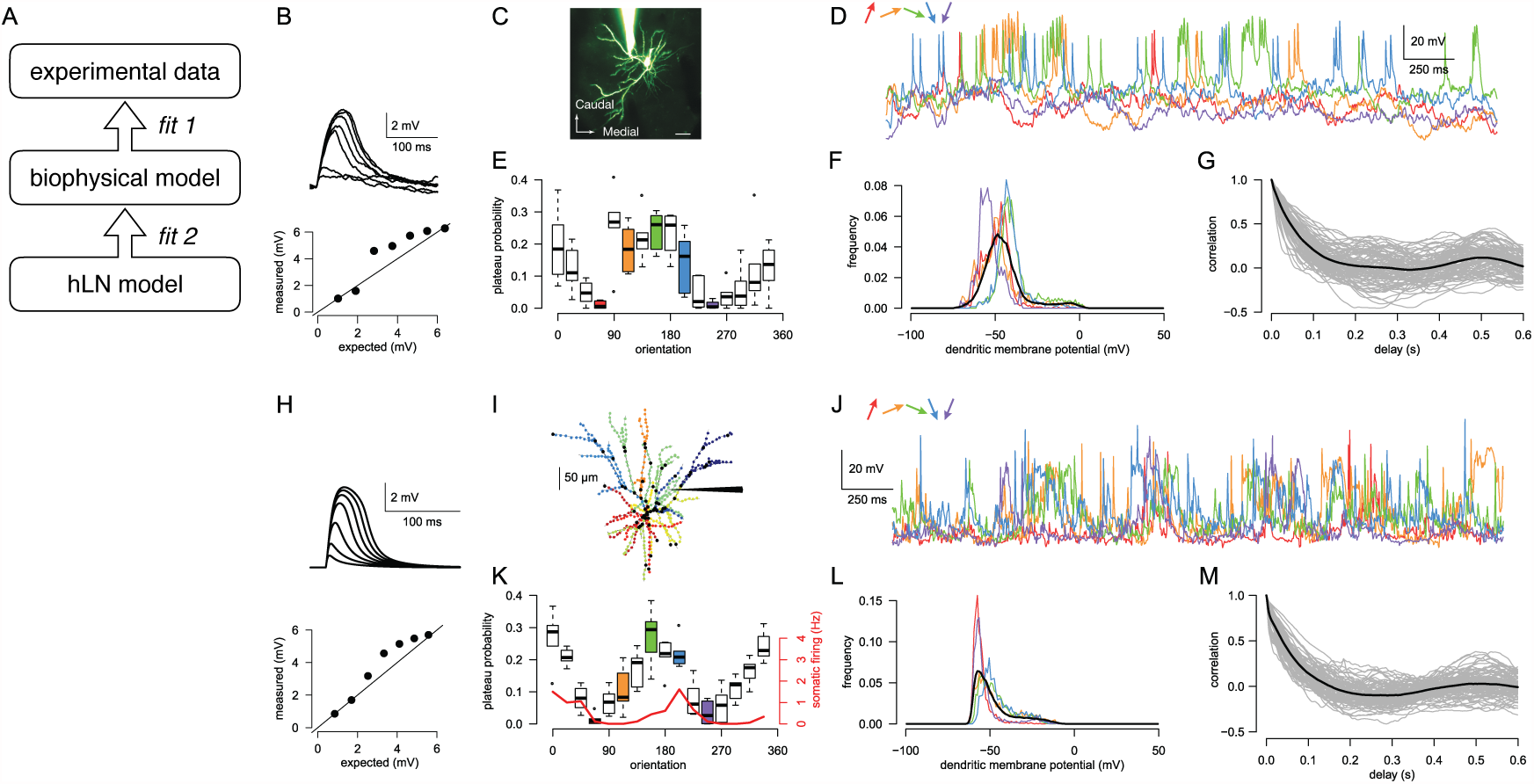
Matching a biophysical model to in *vivo* data. (A) Logic of the approach. We first matched the dendritic and the somatic response of a detailed biophysical model to *in vivo* data (fit 1). This step was required because there are no experimental measurements of the spatio-temporal activation profile for all synapses in a neuron *in vivo*, and its corresponding output. Next, we tuned the parameters of the phenomenological hLN model to match the somatic membrane potential time course of the biophysical model in response to known synaptic inputs (fit 2). (B) Experimental data showing non-linear dendritic integration in layer 2/3 pyramidal neuron *in vitro* (reanalysed from Branco et al. (2010)) (C) Top, two-photon microscopy image (maximum intensity projection) of an Alexa Fluor 594-filled layer 2/3 pyramidal neuron in the mouse visual cortex during a dendritic patch-clamp recording *in vivo* (scale bar, 20 *μ*m). Bottom, examples of membrane potential recordings from a single dendrite in response to differently oriented drifting gratings (colors). Experimental data from Smith et al. (2013). The same dendrite is analysed in E-G. (E) Orientation tuning of the dendritic branch. (F) Histogram of the dendritic membrane potential for different sample input orientations (colors as in D-E) and the average across all different orientations (black). (G) Auto-correlation of the dendritic membrane potential (grey: individual traces for each orientation and repetition, black: average). (H) Non-linear dendritic integration in a biophysical model layer 2/3 pyramidal neuron. (I) The morphology of the reconstructed neuron and the distribution of inhibitory (black dots) and excitatory synapses (colors indicate synapses receiving correlated inputs). Schematic electrode points to dendrite analysed in J-M. (J) Membrane potential of the model recorded in a dendritic branch in response to sustained, *in vivo*-like inputs corresponding to different orientations (colors as in D). (K-M) Orientation tuning (K), and membrane potential histogram (L) and auto-correlation (M) of the dendrite. Colors and symbols are as in E-G, red line in K shows somatic orientation tuning.

We chose layer 2/3 neocortical neurons in the visual cortex because their biophysical properties (Larkum et al., 2007; Branco et al., 2010; Branco & Häusser, 2011) and *in vivo* somatic (Poulet & Petersen, 2008; Smith et al., 2013; Petersen & Crochet, 2013; Polack et al., 2013) and dendritic (Smith et al., 2013; Palmer et al., 2014) activity have been well characterized. To replicate input integration in layer 2/3 neurons under *in vivo*-like conditions, we used a previously validated biophysical model that reproduced dendritic non-linearities observed *in vitro* (Figure 2B,H, Branco et al. 2010; Smith et al. 2013), and tuned the statistics of its input spike trains to reproduce the dendritic and somatic membrane potential dynamics measured experimentally *in vivo*.

During visual stimulation with drifting gratings, the dendritic activity of layer 2/3 cells (Figure 2C, Smith et al. 2013) is characterised by intermittent, large amplitude, plateau-like depolarising events (Figure 2D), reflected in the bimodal dendritic membrane potential distribution (Figure 2F) and slow decay of the autocorrelation function (Figure 2G, see Smith et al. 2013 for experimental details). These plateau potentials reflect the recruitment of dendritic non-linearities (Palmer et al., 2014) and have been shown to enhance the selectivity of orientation tuning (Smith et al., 2013). In line with this, the particular orientation tuning of the dendrite was apparent when we plotted the probability of the plateaus as a function of stimulus orientation for the experimental data from Smith et al. 2013 (Figure 2E), while the weak periodicity of the dendritic activity (second peak of the auto-correlation function, Figure 2G at 0.5 s) could be attributed to the phase preference of dendritic plateaus.

To tune the input statistics in our biophysical model neuron, we chose presynaptic stimulation patterns that reproduced dendritic plateau-potentials and their correlation with visual stimuli of different orientations. To achieve this, we added >600 excitatory (with AMPA and NMDA receptors) and >200 inhibitory synapses (with GABA-A receptors) to the neuron, where the majority of the inhibitory synapses were located near the soma and all other synapses were distributed uniformly throughout the entire dendritic tree (Figure 2I). We found that a relatively low rate of background excitation (5 Hz) and inhibition (20 Hz), alternating with elevated excitatory synaptic activity (20 Hz) balanced by increased synaptic inhibition (Haider et al. 2013, 30 Hz) could reproduce the experimentally observed bimodality of dendritic membrane potentials (Figure 2L).

Although excitation and inhibition were balanced overall, the differential spatial distribution of excitatory and inhibitory synapses caused a disproportionate increase in the dendritic membrane potential during elevated activity, which led to NMDA receptor dependent plateau potentials in distal branches (Figure 2J). The duration of the elevated states was chosen to match the decay of the autocorrelation (Figure 2M), and clustering of co-active presynaptic inputs on the same dendritic branch, as expected (Takahashi et al., 2012; Wilson et al., 2016), facilitated the induction of dendritic plateaus. To model dendritic orientation selectivity, we made the probability of the elevated activity states dependent on both the orientation and the phase of the drifting grating stimulus (Figure 2K; Experimental Procedures), and we subdivided the dendritic tree into thirteen domains that differed in the preferred orientation of their inputs covering a 90° range (Figure 2I). As a result, the soma also showed orientation tuning, such that the peak firing rate of the neuron was between 1 and 2 Hz (Figure 2K, red), matching experimental data (Polack et al., 2013; Smith et al., 2013).

Having extensively validated this biophysical model on experimental data, we next used it as a testbed to analyse dendritic processing under *in vivo*-like conditions.

### A hierarchical linear-nonlinear model of dendritic integration

In order to capture the high-dimensional and potentially complex input-output mapping of a cell under *in vivo*-like conditions, including the effects of non-linear dendritic processing, we adapted a hierarchical extension of the widely used linear-nonlinear (LN) model (Vintch et al., 2015; Freeman et al., 2015). In our hierarchical LN (hLN) model, the input-output transformation of the cell was formalised as a hierarchy of simple subunits (Figure 3A, gray boxes), such that inputs to the same subunit (Figure 3, red and blue spike trains) were first linearly integrated both temporally (using a mixture of standard alpha function synaptic kernels; Figure 3A, orange and purple) and spatially (Figure 3A, yellow), and a separate sigmoidal non-linearity acted on the output of each subunit (Figure 3A, green) before it was linearly combined again with the outputs of other subunits. The form of this model was motivated by previous studies, suggesting that the effects of active dendritic processes can be qualitatively captured by a similar compartmentalisation of dendritic non-linearities into individual branches (Poirazi et al., 2003a; Polsky et al., 2004). Thus, we interpreted the subunits in our model as simplified models of dendritic branches, including the effects of synaptic processing (the kernels) and non-linear input integration (the combination of summation and a static non-linearity).

**Figure 3.**
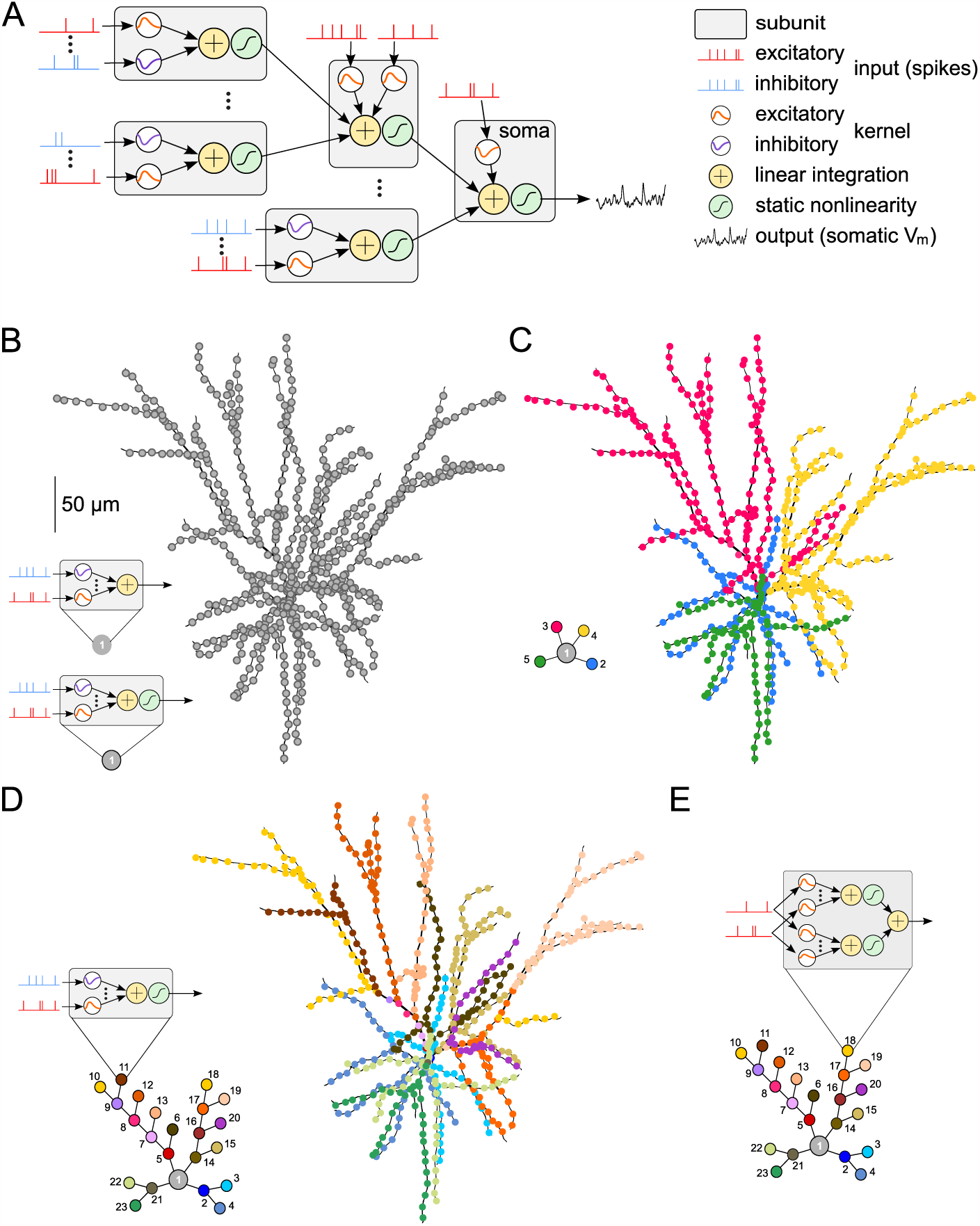
The hierarchical linear-nonlinear (hLN) model. (A) Schematic of an hLN model with four dendritic and one somatic subunit shown. Each subunit (gray boxes) receives input from a number of excitatory (red) and inhibitory spike trains (blue), filtered by positive (orange) or negative synaptic kernels (purple). The filtered inputs are summed linearly (yellow) and passed through a sigmoidal non-linearity (green) before being integrated at the next stage of the hierarchy. (B-E) hLN model architectures of increasing complexity (left) capturing synaptic integration in the biophysical model (right). Each colored circle of an hLN model corresponds to an individual subunit with input spike trains from a subset of synapses (colored dots for excitatory synapses shown on the biophysical model morphology) and an output non-linearity (except for the single subunit of the model in B, top, see also insets). Gray circles correspond to somatic subunits. Models from B to D expand the depth of the hierarchy from a 1-subunit (1-layer) “point neuron” model (B), to a 23-subunit (6-layer) model (D). Model in E expands the breadth of the hierarchy by multiplexing synaptic inputs such that each input spike train feeds into two parallel input channels with different synaptic kernels and non-linearity (inset shows the schematic of a single, multiplexing subunit).

To better dissect the contributions of dendritic non-linearities from those of (peri-)somatic non-linearities responsible for the generation of action potentials, we focussed on the subthreshold somatic membrane potential by setting the density of somatic voltage-dependent sodium channels to zero in the biophysical model (Figure 3A, black trace to the right). (See Figure S4 for results with fitting both the sub- and suprathreshold behavior of the somatic membrane potential.) The parameters of the model (amplitude and time constants of excitatory and inhibitory synaptic kernels, the thresholds of the non-linearities, and the output weight of each subunit) were fit to simultaneously recorded input-output data (input spike trains and somatic membrane potential obtained from the biophysical model) using principled, maximum likelihood-based statistical techniques (Experimental Procedures). We rigorously validated both our statistical methods for model fitting and the ability of the hLN model class to correctly capture the integration of spatially distributed inputs, despite its drastic discretization of the cell’s morphology into a small number of independent subunits (Figures S2-S3).

### Global input-output transformation in pyramidal neurons

We formalised alternative hypotheses about the functional form of dendritic input integration by generating a sequence of increasingly complex architectures within the hLN class that differed in whether their final subunit integrated its inputs purely linearly or it also included a terminal non-linearity (Figure 3B), and in the number of dendritic subunits capable of local non-linear processing (Figure 3B-D). The architectures of these hLN models followed the morphology of the biophysical model and its input distribution as much as possible with the given number of subunits (cf. Figure 2I). We then fitted each of these models to the same data set generated by the biophysical model that we had previously calibrated on *in vivo* experimental data (Figure 2), such that the inputs were the spike trains received by the biophysical model (Figure 4A) and the output was its somatic membrane potential (Figure 4B). We quantified hLN model accuracy by the fraction of variance it explained in cross validation, on a held-out test data set (Figure 4C).

**Figure 4.**
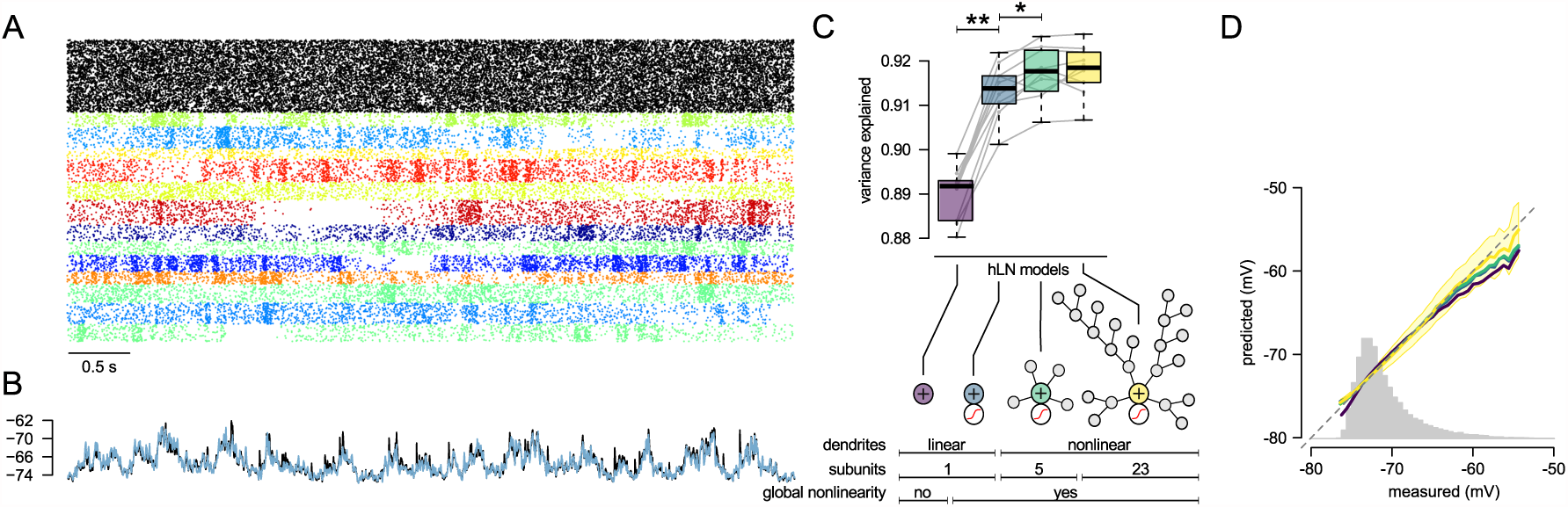
The hLN model class accurately captures dendritic integration. (A) Presynaptic inhibitory (black) and excitatory input spike trains (colors, as in Figure 2I) used for fitting the biophysical model to experimental data (Figure 2). (B) The somatic membrane potential in the biophysical model (black) and predicted by the hLN model with non-linear soma and linear dendrites (blue) in response to the input shown in A. Parameters of the biophysical model and the inputs were identical to that shown in Figure 2, except that somatic active conductances were removed. (C) Prediction accuracy (variance explained) of hLN models with increasing complexity. Bottom shows architecture of different hLN models and table summarizing their main properties (cf. Figure 3B-D). (D) Mean of model predictions as a function of the measured response (colored lines). Gray histogram shows the distribution of the measured response, black diagonal shows identity line. Shaded area indicates the standard deviation of the 23-subunit model’s prediction.

As expected, because the sequence of models was such that more complex models always included the simpler ones as special cases, successful fitting of these models led to monotonically increasing accuracy in predicting the biophysical model’s behavior (Figure 4C). Nevertheless, we found that models including a single subunit, and thus performing linear processing, either without (Figure 3B, top) or with a global non-linearity (Figure 3B, bottom), already explained 90% variance (Figure 4B-C, purple and blue). Introducing more subunits, and hence more localised non-linear processing in the hLN model (Figure 3C-D) increased its accuracy only slightly, but significantly (p<0.001), to 91.5% (Figure 4C, green and yellow). In particular, adding more dendritic subunits corrected the downward bias in predicting high somatic membrane potentials but did not reduce the variance of hLN predictions (Figure 4D).

To test the generality of these findings, we repeated the same analyses by fitting our hLN models to data obtained by stimulating the biophysical model with a wider range of spatially and temporally structured input spike patterns (Figures S5-S6). We systematically varied two parameters of the input patterns that were expected to have a strong influence on the extent to which the cell integrated inputs linearly or non-linearly: input firing rates and the number of presynaptic ensembles organised into functional clusters (groups of synapses with correlated inputs). In all cases tested we found that linear models accounted for at least 80% of variance, and for at least 90% within the physiologically feasible regime (maximal input rates above 10 Hz, and ensemble number above 1 and below 104), with even multiple layers of local non-linearities only improving predictions by at most 2%. Taken together, these results suggest that within the parameter space that we have explored, local non-linear processing in dendrites does not account for more than 10-20% of the somatic membrane potential variance, and that the input-output transformation performed by non-linear dendrites can be well described by linear processes followed by a global non-linearity.

The relatively modest extent, ~10%, to which the input-output transformation in dendrites was found to be nonlinear does not indicate that active conductances have a negligible contribution to input integration (Cash & Yuste, 1999). To study how they influence these linear transformations, we compared hLN models with linear dendrites (Figure 4C, blue) that were fit either to the original biophysical model (Figure 5A, black), including a rich repertoire of active conductances (Methods), or to a passive variant (Figure 5A, gray) which was identical to the original model in its morphology, passive membrane properties, and the input spike trains it received, but all active conductances (including NMDA receptors) were absent from it. We found that, as expected, the linear model provided a slightly better fit to the passive than to the active cell (Figure 5B, blue, 95% vs. 91.5%, p<0.001). More crucially however, although the difference in the quality of fits was small, the synaptic kernels underlying these fits were drastically different. In particular, excitatory kernels were larger and longer-lasting in active dendrites (Figure 5C, top, D-E, orange), mainly reflecting the recruitment of NMDA-receptor currents, while inhibitory kernels became larger (reflecting the an increase in driving force caused by larger overall dendritic depolarization) but remained similar in their time courses (Figure 5C, bottom, D-E, purple). Thus, beside turning the input-output transformation slightly nonlinear, the primary effect of the active dendritic conductances is to change the linear integration properties of the cell.

**Figure 5.**
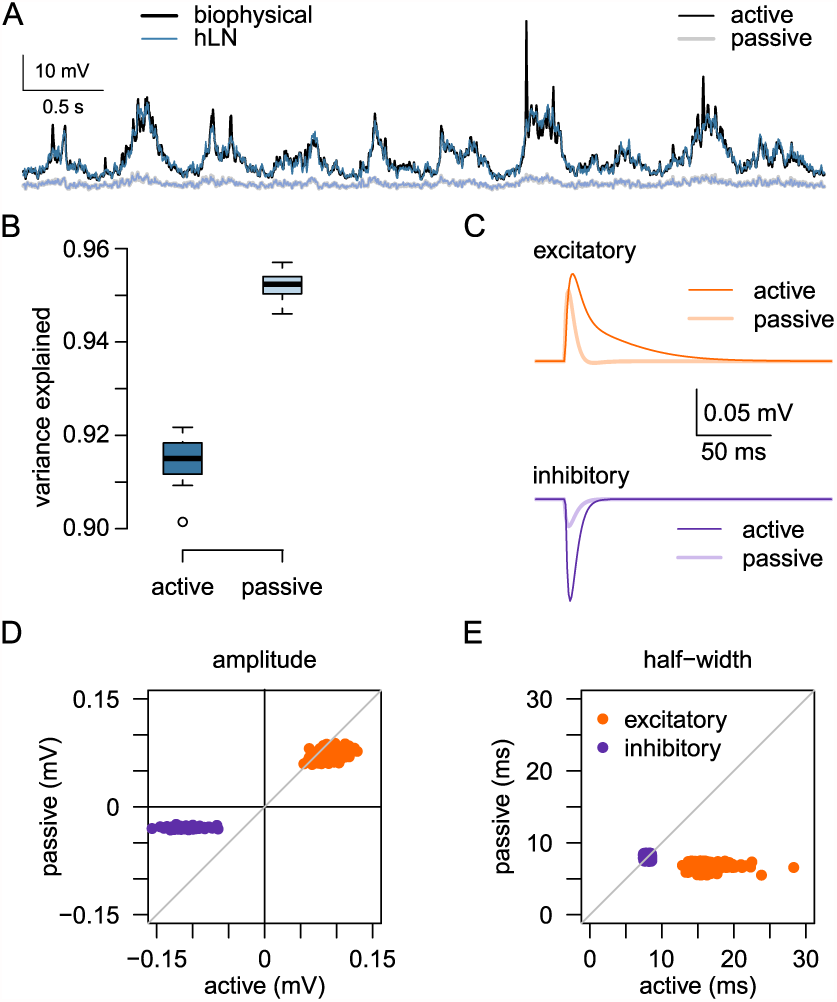
Effects of active dendritic conductances on the properties of linear input-output transformation in dendrites. (A) Somatic membrane potential in the active (black) and passive (gray) biophysical neuron model to *in vivo*-like stimulation (as in Figures 2 and 4) together with the hLN model’s prediction (dark and light blue, respectively). (B) Variance explained by the hLN model for the active (dark blue) and passive cell (light blue). (C) Average excitatory (top, orange) and inhibitory synaptic kernels (bottom, purple) for fitting the responses of the active (dark colors) or passive cell (light colors). (D-E) Amplitude (D) and half-width (E) of individual excitatory (orange dots) and inhibitory synaptic kernels (purple dots) for fitting the active vs. the passive model. Gray diagonals show identity.

### Input multiplexing

Formalising the input-output mapping of single neurons in terms of a hierarchical linear-nonlinear model allowed us to identify functional architectures that had not been considered previously (Häusser & Mel, 2003) but which follow logically from this model class and therefore may present viable descriptions of the effects of non-linear dendrites. In particular, while the transition from a single layer (single subunit) to multilayered (multiple subunits) hierarchies represents an increase in *depth* of the architecture, we wondered whether increasing its *breadth* may also increase its predictive power. To test this, we multiplexed each input spike train to each subunit into two channels, thus allowing the integration of inputs independently by kernels with potentially different parameters (Figure 3E).

We found that multiplexing could substantially improve model accuracy (Figure 6A). In particular, the advantage of multiplexed architectures over non-multiplexed ones (Figure 6A, orange vs. yellow) was greater than that of going from a single to 23 subunits (Figure 6A, yellow vs. blue). In order to understand what aspects of neuronal input integration were captured by multiplexing, we considered a simplified case, in which only four dendrites were stimulated (Figure 6B) with patterns similar to those producing realistic dendritic membrane potential distributions (Figure 4) and autocorrelograms (Figure 6C). We then compared the best-fit parameters of the two input channels of individual subunits (Figure 6D-H) and found a robust separation of input channels for their ‘speed’, i.e. the time constant with which they integrate excitatory inputs: 5.9 ± 1.4 ms vs. 26.2 ± 3.2 ms (Figure 6D, E). The fast and slow channels also differed in the balance of excitatory vs. inhibitory inputs they integrated, with inhibitory synapses contributing much more to the slow than the fast channel (Figure 6D, F). Moreover, the threshold of the sigmoidal non-linearity was also different in the two channels, such that most of the input distribution lay on the supralinear-linear vs. linear-sublinear part of the sigmoidal non-linearity in the slow and the fast channels, respectively (Figure 6D). This resulted in the slower channel applying higher average gain (Figure 6G) and a more strongly supralinear transformation to its inputs than the fast channel (Figure 6H).

**Figure 6.**
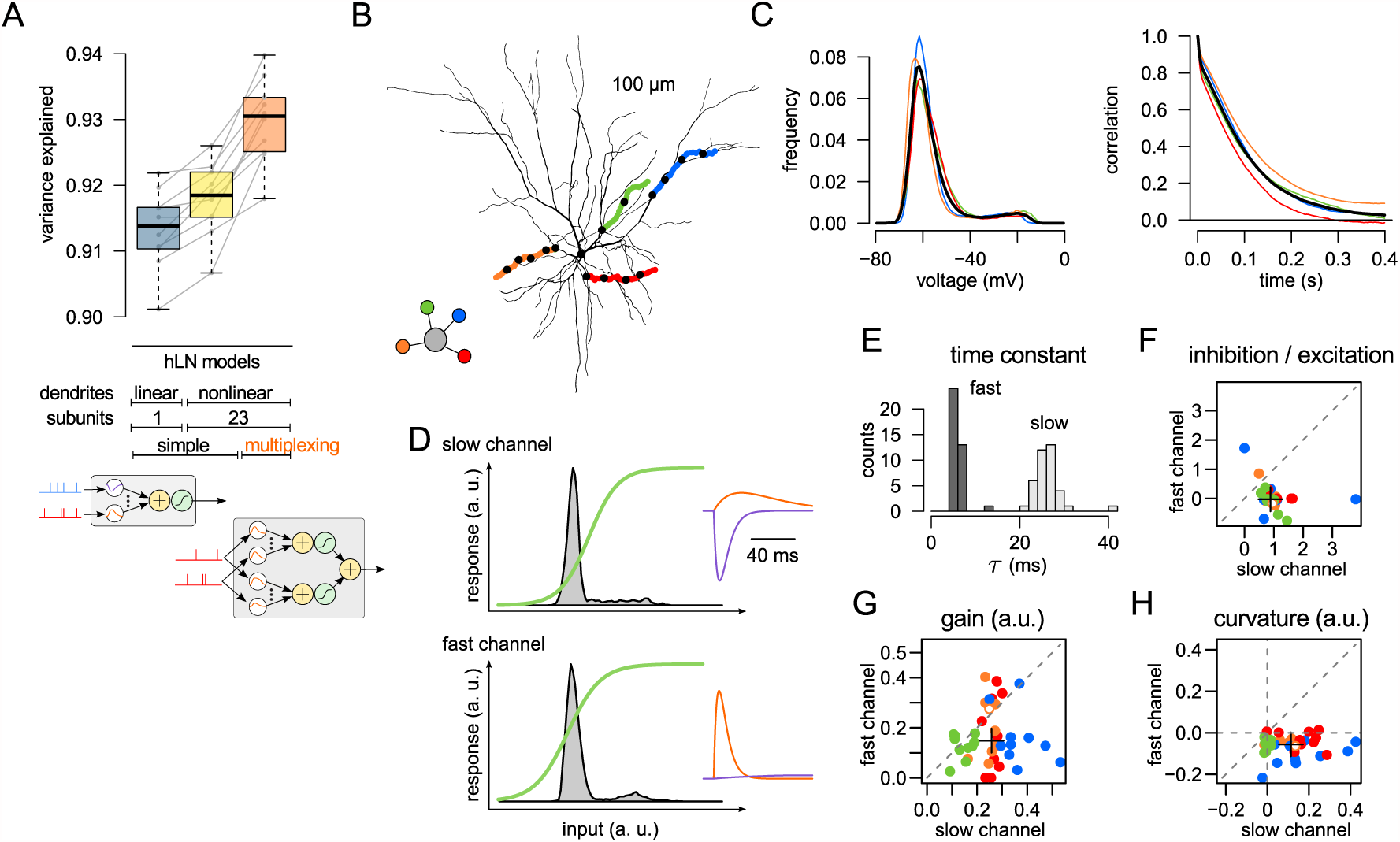
Input multiplexing. (A) Prediction accuracy (variance explained) of hLN models with increasing complexity, including multiplexing (orange). Table in middle summarizes the main properties of different hLN models (cf. Figure 4C), bottom illustrates difference between non-multiplexing (left) and multiplexing subunits (right, cf. Figure 3D-E). Blue, yellow: same as in Figure 4C, shown here for reference. (B) Biophysical cell model with four dendrites stimulated (colored) and the architecture of the hLN model fitted to its responses (inset). (C) Membrane potential distributions (left) and autocorrelaograms (right) in individual dendrites (colors as in B) and their average (black). (D) Properties of the two input channels (top: slow, bottom: fast) in a representative subunit. Gray histograms: distributions of excitatory synaptic inputs after temporal filtering with the corresponding synaptic kernels, green lines: output non-linearities of the input channels. Insets show excitatory (orange) and inhibitory synaptic kernels (purple). (E) Distribution of excitatory time constants in the two input channels (dark vs. light gray) across the four subunits and 10 independent fits. (F-H) Ratio of inhibitory to excitatory synaptic kernel amplitudes (F), and the slope (G) and curvature (H) of the output nonlinearity (averaged under the filtered input distribution, see gray histograms in D) in fast vs. slow input channels across subunits (colors as in B) and 10 independent fits. Negative or positive curvature in H implies sublinear or superlinear integration, respectively. Crosses in F-H indicate population medians, empty circles correspond to the example shown in D.

In sum, the fast (more sublinear) input channel exhibited AMPA channel-like properties, while the slow (more supra-linear) channel combined properties of NMDA channel-and GABA-mediated transmission. The incorporation of inhibition into the slow input channel reflects the increased driving force for the GABAergic synapses during periods of larger depolarisation, which are well captured by the slow input channel because of its supra-linearity and higher overall gain. Integrating inhibition with the fast channel, where a large fraction of input is processed sublinearly, would have resulted in a smaller effect of inhibition at higher levels of depolarisation, and separating inhibition to an altogether separate input channel would have resulted in a simple linear integration of excitatory and inhibitory inputs, with the effect of inhibition being independent of the level of depolarisation.

### A case study: dendritic integration in cerebellar granule cells

The main advantage of our method is that it allows the joint estimation of all parameters of the functional architecture of input-output transformations in a cell (kernels, non-linearities, and their hierarchy) during *in vivo*-like conditions, without the need to conduct piecemeal minimal stimulation experiments and simulations, and analyses that may ultimately not generalise well to the *in vivo* case (Figure 1). However, as the ground truth for the functional architecture of a neuron is generally unknown, and may in fact lie outside any particular model class considered (in our case, the hLN model), the validity of our results can only be measured by simple summary statistics, such as variance explained (Figures 4-6). In order to conduct a more stringent test, we applied our analysis to cerebellar granule cells, whose input and output has been well characterised *in vivo* and whose functional architecture is relatively well understood (Figure 7). In particular, granule cells are electrotonically compact, which means that their dendrites do not effectively compartmentalise their non-linearities and thus are unlikely to implement multilayered functional architectures.

**Figure 7.**
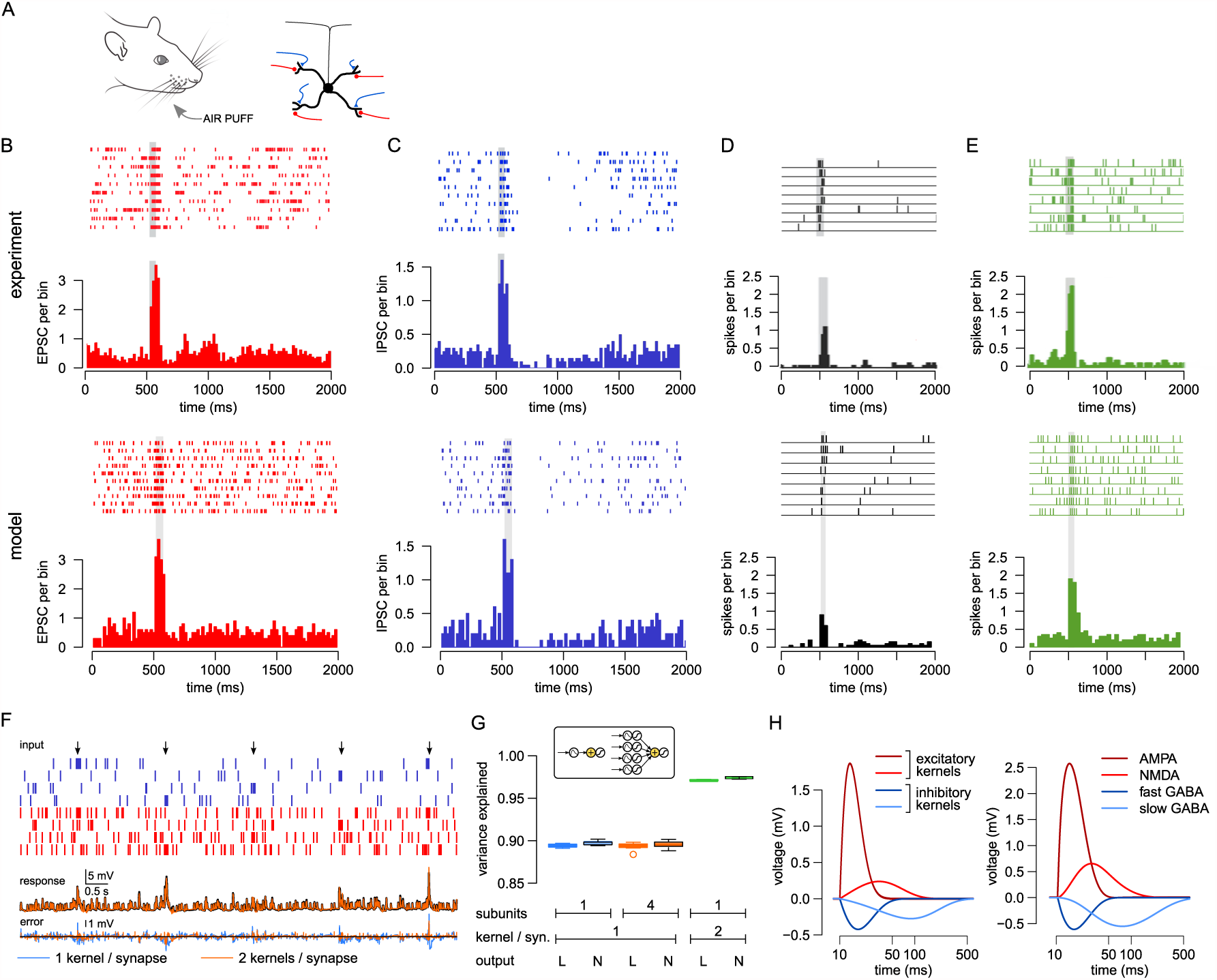
Dendritic integration in cerebellar granule cells *in vivo*. (A) Schematics of the experimental configuration (left) and the granule cell morphology, with excitatory (red) and inhibitory (blue) synaptic contacts made on its four dendritic claws (right). (B-E) Matching the inputs (B-C) and the output (D-E) of the biophysical model neuron (bottom) to experimental data (top) from Duguid et al. (2015) (B-C) and Duguid et al. (2012) (D-E). B: Raster (ticks) and peri-stimulus time histogram of input EPSPs recorded *in vivo* from cerebellar granule cells (top) together with the synaptic inputs used in the model (bottom). C: IPSPs *in vivo* (top) and in the model (bottom). D: Somatic action potentials under control input conditions recorded *in vivo* (top) and in the model (bottom). E: similar to D but after blocking GABA_A_ inhibitory inputs with gabazine. Shaded area in B-E indicates period of stimulus presentation. (F) Top: *in vivo*-like synaptic input patterns. Each of the four dendrites recives inputs from a pair of excitatory (red) and inhibitory synapses (blue) with correlated activity. Arrows indicate sensory stimulation. Middle: the subthreshold membrane potential response of the biophysical model (black), and the prediction of the linear hLN model with a mixture of kernels (orange). Bottom: error between the response of the biophysical model and the prediction of the linear model with simple (blue) or mixture kernels (orange). (G) Prediction accuracy of hLN models. Insets show model hLN architectures (only excitatory inputs are shown for clarity). Models with simple kernels (blue and green) have one inhibitory and one excitatory synaptic kernel in each subunit. The multi-layered model (green) has four non-linear subunits each corresponding to one dendritic branch of the biophysical model. The model with mixture kernels (orange) integrates both excitatory and inhibitory inputs with two different synaptic kernels. Each model has two variants: with (box plots with black contour) or without a somatic non-linearity (box plots without black contour). (H-I) Synaptic kernels recovered from the hLN model by fitting *in vivo*-like input-output mapping (left) and individual synaptic responses in the biophysical model (right). Note logarithmic time axis.

Similarly to our approach to L2/3 neurons, we started by constructing a biophysically detailed model of a granule cell (Figure 7A, right) which we validated against *in vivo* experimental recordings during the application of brief air puffs to the whiskers and the peri-oral surface (Figure 7A, left; Duguid et al. 2012, 2015). This model reproduced the statistics of both the excitatory and inhibitory inputs (Figure 7B-C) as well as the output firing rate dynamics of the cell during control conditions (Figure 7D) and the application of the GABAA receptor antagonist gabazine (Figure 7E). We then stimulated this biophysical model cell with *in vivo*-like input patterns (Figure 7F, top) and fitted hLN models to its subtreshold somatic membrane potential (Figure 7F, middle, black).

We found that a simple linear model accounted for 90% of variance (Figure 7F, bottom, blue, and G, blue), with a global non-linearity adding little improvement to the accuracy (Figure 7G, blue with black outline). Adding further layers of non-linear subunits also did not improve accuracy (Figure 7G, orange). In contrast, introducing a mixture of synaptic kernels for both excitatory and inhibitory inputs improved accuracy to 97% (Figure 7F, middle and bottom, orange, and G, orange), with an additional global non-linearity again contributing little (Figure 7G, orange with black outline). Moreover, the four functional synaptic kernels (two excitatory and two inhibitory) recovered by the best-fit model with a mixture of kernels (Figure 7H, left) closely resembled the individual PSPs corresponding to the four different receptor channels (AMPA and NMDA for excitation, and fast and slow GABA for inhibition) in the biophysical model (Figure 7H, right). Quantitatively, the estimated kernels of the hLN model fitting *in vivo*-like data were somewhat smaller in amplitude and faster than the PSPs in the biophysical model because the increased total membrane conductance during *in vivo*-like conditions rendered membrane potential dynamics more leaky, thus decreasing both the input resistance and time constant of the cell. Taken together, these results confirm that our method is able to recover individual synaptic contributions to dendritic integration during *in vivo*-like conditions and, more specifically, that dendritic integration in granule cells is dominated by linear processing with a mixture of time scales rather than non-linear hierarchical processing.

## Discussion

We have introduced a novel model-based approach for analyzing dendritic integration and describing the inputoutput transformation of single neurons with complex dendritic trees receiving *in vivo*-like input patterns. The main advances of this work are the development of the new analysis methods and the unexpected finding that synaptic integration in non-linear dendrites can be well described by linear filters and a single, global non-linearity - that is, without multiple layers of local non-linearities suggested by earlier work (Poirazi et al., 2003b). For this approach, we developed a flexible and powerful model class, the hLN model, that provides a compact mathematical characterization of the input-output transformations of individual neurons and can be efficiently fitted to data. We applied our approach to analyze the integration of direction-selective inputs in a biophysical model of a L2/3 pyramidal neuron, and found that a very large fraction of the neuronal response is successfully captured by linear integration. Moreover, our analysis also showed that multiplexing the inputs to *parallel* (slow and fast) processing channels within computational subunits is a form of non-linear dendritic processing that can be equally important as the classically considered *serial* hierarchy of subunits (Häusser & Mel, 2003). We demonstrated the generality of this approach by applying it to a different cell type, cerebellar granule cells, which revealed fundamentally linear integration while fully capturing the different excitatory and inhibitory synaptic time scales that have been recorded *in vivo*.

### Synaptic integration under *in* vivo-like input conditions

Determining the input-output transformations implemented by single neurons under *in vivo*-like conditions is a key step towards understanding the role of single neurons in behaviourally relevant computations. Neurons have complex and diverse morphology and physiology, but early theoretical work mostly ignored this complexity and reduced neurons to single point integrators (McCulloch & Pitts, 1943). Subsequent studies aimed to characterize neurons as complex, non-linear dynamical systems (Hodgkin, 1948; Izhikevich, 2007), as well as explicitly modelling synaptic integration of inputs on spatially extended dendritic branches (Rall, 1962). Rall’s work in particular demonstrated the critical influence of the cable properties of dendrites on synaptic integration, and paved the way for many theoretical and experimental studies on dendritic integration (Segev & Rall, 1998). However, because of technical limitations, the majority of these studies evaluated integration of input patterns that are substantially simpler than the spatio-temporally complex spike patterns characteristic of *in vivo* population activity (Cunningham & Yu, 2014). For example, direct current injection in dendrites (Koch, 1999; Izhikevich, 2007), electrical synaptic stimulation (Schiller et al., 2000) or glutamate uncaging (Losonczy & Magee, 2006) are all typically limited to small regions of the dendritic tree and to temporal patterns of limited dimensionality, and while extremely useful for characterizing the basic integrative properties of single neurons, it remains unclear how such properties are engaged by complex input patterns. In particular, non-linear systems can be exquisitely sensitive to input conditions (Strogatz, 1994) and thus to gain insight into the neuronal input-output transformations under *in vivo* conditions it is important to evaluate single neuron responses to *in vivo*-like stimuli.

Previous theoretical work on neuronal input processing during *in vivo*-like conditions has mainly focused on the increase in input conductance caused by persistent synaptic bombardment (the ‘high conductance state’), and analyzed its effects on the efficacy of synaptic inputs (Destexhe et al., 2003) and on events such as the initiation and propagation of dendritic spikes (Rudolph & Destexhe, 2003; Williams, 2004; Jarsky et al., 2005; Farinella et al., 2014). While these studies highlighted important differences in synaptic integration between the quiescent and the *in vivo*-like states, and provided a means to evaluate the processing of complex input patterns, they did not describe the input-output transformation of individual neurons during *in vivo*-like input conditions. The approach we have developed provides a principled way of achieving this and can be applied to data from both compartmental models and experiments. In the present paper we applied this technique to analyze synaptic input processing in a multi-compartmental biophysical model of a cortical neuron endowed with several active dendritic mechanisms. The model parameters were carefully matched to experimental data, and while no model can be a perfect replica of the original system, we found that our conclusions are robust against changes in the input firing rate or the degree of synaptic clustering, even beyond their biologically realistic range (Figs. S5-S6). Moreover, we believe that our methods pave the way for similar analyses of real neurons, once simultaneous recordings of their input spike trains and somatic membrane potential become available. (Grienberger & Konnerth, 2012).

### Compact mathematical characterization

Developing compact mathematical characterizations of the inputoutput transformations of individual neurons is a long-standing challenge (Gerstner & Naud, 2009; Poirazi et al., 2003b) that is a critical step for understanding the population dynamics and computations that emerge at the level of neural circuits (Ahmadian et al., 2013; Rubin et al., 2015). However, classical principled methods for distilling simplified single neuron models are only formally valid for electrotonically compact neurons (Koch, 1999), in which the contribution of dendritic processes for synaptic integration is minimal, and for neurons with passive dendrites that lack voltage-dependent conductances (Rall, 1962) (but see also Marasco et al. 2012; Rossert et al. 2017). Similarly, due to the vast complexity of dendritic non-linearities and the current lack of a formalization of their contributions to single neuron computations, the majority of theories of network-level computations either rely on single-compartmental models (Dayan & Abbott, 2001), thereby ignoring the role of dendrites, assume linear dendritic processing (Cook & Johnston, 1997), or make very specific assumptions about the form of dendritic nonlinearities based on largely qualitative arguments (eg. about coincidence detection (Pecevski et al., 2011; Kaifosh & Losonczy, 2016)). Our framework offers a principled way of estimating the contribution of non-linear dendritic processing to the response of neurons and incorporating it efficiently in single neuron models designed for network simulations.

### hLN models

To address the challenges raised by the high dimension of inputs with a complex spatio-temporal structure, we formalised the mapping from input spikes to the somatic membrane potential using a hierarchical parametric model that could be efficiently fitted to data. This approach enabled us to compare several hypotheses about the form of the input-output transformation in single cortical neurons by varying the number and arrangement of model subunits and, after fitting each configuration to data, comparing their accuracy in predicting real responses (on a held-out test data set). Importantly, this model comparison allowed us to measure the impact of different factors, such as dendritic non-linearities, on the overall input-output transformation of the cell. Our framework is flexible: further configurations could be designed to match the non-linearities exhibited by different cell types or test alternative hypothesis about the form of signal transformation in pyramidal neurons under specific input conditions.

Our approach has its roots in system identification (Wu et al., 2006) applied to modelling of visual signal processing in the retina (Meister & Berry, 1999; Schwartz et al., 2006) or of neuronal responses to somatic and dendritic current injections (Poliakov et al., 1997; Cook et al., 2007). While the focus of system identification in systems neuroscience has mainly been on the mapping between an *analog* stimulus (e.g.: visual pixel intensities or input currents) and the *digital* spiking response of the recorded neuron (Schwartz et al., 2006), the approach we have developed reverses these classes by deriving a mapping between the digital presynaptic spike trains and the analog somatic membrane potential.

In recent work, models aiming at predicting spike timings have used spike trains of the same or different neurons to capture adaptation, as well as the effect of inter-neuronal functional couplings (Pillow et al., 2008; Truccolo et al., 2010). However, in these studies the stimuli used for predicting neuronal responses are not the stimuli directly received by the cell, and therefore cannot be used to identify the contributions of single neuron integration to signal processing, and separate them from network processes. The recognition of this shortcoming led to the development of various hierarchical extensions of the LN framework (McFarland et al., 2013; Goris et al., 2015; Vintch et al., 2015; Freeman et al., 2015; Maheswaranathan et al., 2016), albeit these models have been previously applied only in the analog stimulus domain and made specific assumptions about the stimulus structure (e.g., Gaussianity) that are not met by the synaptic inputs used here. Moreover, our model is fitted to the somatic membrane potential of the cell because parameter estimation is more efficient when it is based on the subthreshold voltage than on the spikes (Ramirez et al., 2014) and it better disects the contribution of dendritic nonlinearities to the overall response of the cell. A key advance of our work is that the use of the unique combination of these analysis features enabled us to identify the input-output transformation of single neurons with realistic morphology and synaptic inputs.

It is important to note that the hLN model framework is complementary to biophysical modelling of single neurons. hLN models provide a compact and intuitive description of the input-output transformation implemented by the neuron, but they lack mechanistic insight. In contrast, biophysical models can reveal the physiological processes underlying signal integration and propagation in neurons, but the overall picture of how the neuron transforms information is often hard to grasp (Herz et al., 2006). Moreover, biophysical models accurately matched to data have the potential to generalize across different input conditions (Druckmann et al., 2011), but parameter tuning in these models is challenging (Huys et al., 2006; Friedrich et al., 2014) and simulations are computationally very expensive (Markram et al., 2015). The accuracy of hLN models on the other hand is limited to the specific input conditions that they were trained on (Cook et al., 2007), but they are very efficient when fitted to data and simulated, allowing integration in large scale network simulations.

### Linear integration

Several *in vitro* experimental studies have demonstrated the presence of both supra- and sublinear dendritic processing via a variety of mechanisms, including dendritic Na^+^ (Golding & Spruston, 1998), Ca^2+^ (Schiller et al., 1997; Larkum et al., 1999) and NMDA-receptor spikes (Schiller et al., 2000; Branco et al., 2010; Makara & Magee, 2013), as well as synaptic saturation (Vervaeke et al., 2012; Abrahamsson et al., 2012) and activation of voltage-gated potassium channels (Hoffman et al., 1997; Urban & Barrionuevo, 1998). More recently, many of these dendritic non-linear mechanisms have been recorded *in vivo* and found to participate in computations such as orientation selectivity (Smith et al., 2013), angular tuning (Lavzin et al., 2012) or coincidence detection (Xu et al., 2012), thereby strongly suggesting that dendritic processing is a fundamental component of input processing in neural circuits. In principle, dendritic non-linearities can either substantially change the form of neuronal input integration (Poirazi et al., 2003b) or simply change the gain of linear integration (Cash & Yuste, 1998). The dominant effect likely depends on the synaptic input patterns and mechanisms involved, and determining this requires formalization of the single neuron input-output transformation and evaluation of how it changes under different conditions, such as with varying input statistics or block of specific mechanisms. The approach we have presented here provides a set of analysis tools for achieving this.

By evaluating the computations performed by a biophysical model of a cortical neuron receiving *in vivo*-like inputs, we showed that only 10% of the response variance can be attributed to local non-linear dendritic processes when the input statistics produced membrane potential profiles that matched *in vivo* recordings (Smith et al., 2013). This is consistent with previous estimates of the contribution of non-linear processing to the somatic membrane potential responses of cortical cells (Jolivet et al., 2006; Mensi et al., 2012; Cook et al., 2007; Rossert et al., 2017), and in agreement with the finding that linear processing of input spike counts accounts for ∼80% of the variance in the mean neuronal firing rate (Poirazi et al., 2003b). However, these previous results were based on simple input patterns, such as constant current injection (Jolivet et al., 2006; Mensi et al., 2012; Cook et al., 2007) or constant input firing rates (Poirazi et al., 2003b), or model inputs which were not calibrated on intracellular recordings (and e.g. had almost no correlations) (Rossert et al., 2017), both of which may lead to an underestimation of the effect of nonlinearities (Figure S1, Figure S6). In contrast, the major novelty of our results compared to these previous studies is that we estimated the contribution of dendritic nonlinearities under *in vivo*-like input conditions, after carefully calibrating our inputs to intracellular recordings (Figure 2).

We found that within the parameter space explored in our biophysical model, input integration can be well described by a linear process. This suggests that when inputs are distributed across the dendritic tree, the contribution of local non-linearities to the neuronal input-output transformation is limited, and that dendritic mechanisms mainly act to change the gain of the transformation. Importantly, adding a single, global non-linearity significantly increased the accuracy of our hLN model, more than the subsequent addition of more sub-units and local nonlinearities. This could represent a global dendritic non-linearity, as recently observed experimentally in L5 and CA1 pyramidal neurons (Bittner et al., 2015; Takahashi et al., 2016), which could be a major mode of dendritic computation *in vivo*. It is important to note that in our biophysical model, the dominant non-linearity is current flowing through NMDA receptors (NMDARs), which can exhibit different input amplification regimes (Schiller & Schiller, 2001) and produce graded amplification of inputs (Branco & Häusser, 2011). This property makes NMDARs particularly suited to control the gain of synaptic integration and therefore be captured by linear integration processes, and the recovered synaptic kernels clearly reflect the recruitment of NMDARs in the active biophysical model.

The apparent linearity of the integration *in vivo* is also supported by the findings that linear models can accurately predict both the spike trains of cortical pyramidal neurons based on the spiking history of other cortical cells (Truccolo et al., 2010) and the neuronal and behavioural response to optical stimulation of various intensity and duration (Histed & Maunsell, 2014). However, the actual inputs the neurons receive was not known in earlier studies thus leaving open the possibility that more clustered inputs could potentially elicit more nonlinear responses. Here we have shown that input integration is approximately linear even when synaptic inputs are clustered (Fig. 4, Fig. S6).

### Clustering and input statistics

Synaptic input clustered in space and time is maximally efficient at activating dendritic non-linearities (Polsky et al., 2004; Makara & Magee, 2013), including NMDARs, especially if synapses are located in high impedance regions of the dendritic tree (Major et al., 2013; Branco & Häusser, 2011). Our simulations aimed to reproduce population activity in the visual cortex of rodents during drifting grating stimulation, which is dominated by low-dimensional cell assembly dynamics where groups of orientation-tuned neurons are transiently and synchronously active (Carrillo-Reid et al., 2015). We modelled presynaptic cell assembly dynamics by strongly correlated firing rate fluctuations (Macke et al., 2011; Ujfalussy et al., 2015), and as discussed above, this generated a regime of NMDAR activation that was compatible with gain modulation and well described by linear processes. However, when we increased the amount of clustering, we found that, as expected, the variance attributed to local non-linearities increased, up to 20%, most likely due to pushing NMDARs towards the spike-generation regime (Major et al., 2013). Although this regime did not agree with *in vivo* dendritic recordings, these results more generally demonstrated the power and versatility of our approach for formalizing and comparing neuronal input-output transformations under a variety of conditions.

The strong negative correlation between the frequency of the dendritic Na^+^-spikes and the hLN-models’ performance (Figure S6J) indicates that most of the unexplained variance arises from dendritically evoked Na+-spikes appearing as spikelets of variable amplitudes in the soma (Golding & Spruston, 1998; Crochet et al., 2004; Smith et al., 2013; Schmidt-Hieber et al., 2017). While in cortical pyramidal cells NMDA receptor activation has been shown to be the primary influence on neuronal responses *in vivo* (Lavzin et al., 2012; Smith et al., 2013; Palmer et al., 2014; Schmidt-Hieber et al., 2017), which our hLN model captures accurately, future developments of the hLN approach could improve its ability to capture events such as the initiation and the propagation of dendritic Na+-spikes. This could be done, for example, by extending the static non-linearities employed here with simplified dynamical models of spike generation and propagation along the network of the hierarchical subunits as proposed recently to model spike responses of neurons with dendritic calcium spikes (Naud et al., 2014). Interestingly, our results also suggest that Na^+^- and NMDA-related non-linearities may involve drastically different hierarchical processing within the dendritic tree, akin to multiplexing inputs to a slow channel with a single global non-linearity and a fast channel with local non-linear interactions and multiple hierarchical steps.

### Input multiplexing

In addition to estimating the contribution of non-linear dendritic processing to the somatic membrane potential, our approach also revealed a novel way of conceptualizing synaptic integration: multiplexing inputs into parallel processing channels with different linear and non-linear integration properties. Similar multiplexing architectures have been previously applied to model input processing by separate subnetworks during phase invariant neuronal responses of complex cells in the visual system (Adelson & Bergen, 1985; Rust et al., 2005; Vintch et al., 2015). This form of non-linear input-integration, which increased the accuracy of our model predictions significantly, represents a major transition from conventional static dendritic non-linearities. In both the pyramidal and granule cell models evaluated here, the main difference between the synaptic kernels was the time scale of input integration, with inputs being processed in parallel by a fast and a slow channel. Importantly, in the case of the pyramidal neuron, the two processing channels targeted different non-linear subunits enabling the neuron to dynamically adjust the properties of integration depending on the input statistics. At low input rates the fast channel has a high gain and therefore its output dominates the neuronal response, while the contribution of the slow channel is relatively small. Conversely, at higher input rates the gain of the supralinearity of the slow channel increases and dominates the response, whereas the fast channel becomes saturated. This arrangement significantly improves the ability of the hLN model to capture the dynamics of NMDAR-dependent integration. Moreover, mutliplexing inhibitory inputs resulted in most inhibition being processed via the slow, super-linear channel, which reflects the increased depolarization and consequently increased driving force for GABAergic currents during the engagement of the slow excitatory channel. These results demonstrate the ability of multiplexing to capture important biophysical effects, and suggest that this approach will be useful for abstracting the compound effects of multiple conductances with different dynamic properties, without having to model them explicitly. The application of the hLN mutliplexing motif to cerebellar granule cells further underscored the flexibility and generality of our approach, as the hLN model not only captured synaptic integration with high accuracy, but also recovered kernels that had direct correlates with synaptic mechanisms observed *in vivo*.

## Methods

### Biophysical models

Simulations were performed with the NEURON simulation environment (version 7.4) using a detailed reconstruction of a biocytin-filled layer 2 pyramidal neuron (NeuroMorpho.org ID Martin, NMO-00904) as described previously (Smith et al., 2013). Briefly, the passive parameters were *C_m_* = 1 *μ*F/cm^2^, *R_m_* = 7000 Vcm^2^, *R_i_* = 100 V cm, yielding a somatic input resistance of 70MΩ.

Active conductances were added to all dendritic compartments and occasionally to the soma (Figure 2, Figure S4C) and included the following: voltage-activated Na^+^ channels (soma 100 mS/cm^2^, dendrite 8 mS/cm^2^ and hot spots 60 mS/cm^2^, Nevian et al. 2007); voltage-activated K^+^ channels (10mS/cm^2^ soma and 0.3 mS/cm^2^ dendrite); M-type K^+^ channels (soma 0.22mS/cm^2^ and dendrite 0.1 mS/cm^2^); Ca^2+^-activated K^+^ channels (soma 0.3 mS/cm^2^ and dendrite 0.3 mS/cm^2^); high-voltage activated Ca^2+^ channels (soma 0.05 mS/cm^2^ and dendrite 0.05 mS/cm^2^) and low-voltage activated Ca^2+^ channels (soma 0.3 mS/cm^2^ and dendrite 0.15 mS/cm^2^). For the simulations shown in Figure 6B-H all active currents were excluded except NMDA receptors.

AMPA, NMDA and GABA-A synapses were modelled as a bi-exponential function, with time constants of AMPA τ_1_ = 0.1 ms, τ_2_ = 2 ms; NMDA τ_1_ = 3ms, τ_2_ = 40ms and GABA-A τ_1_ = 0.1 ms, τ_2_ = 4 ms (the inhibitory reversal potential was set to −80mV). The Mg^2+^ block of NMDA synapses was modelled according to Jahr & Stevens (1993). The kinetic NMDA receptor model used in Figure 1 was modelled after Kampa et al. (2004) and included five states (unbound, closed, open, slow and fast desensitization states). To facilitate the comparison with the results using non-kinetic NMDA receptors we assumed that the Mg^2+^ block of NMDA synapses is instantaneous and was modelled according to Jahr & Stevens (1993).

AMPA and NMDA components were colocalized and coactivated at each excitatory synapse. The maximal conductance of the NMDA synapses were set to g_max_ = 0.5 nS and the AMPA synapses were set to g_max_ = 0.25 nS except in Figure 1 where it was varied between g_max_ = 0.25 nS (default, NAR=2) and g_max_ = 0.75 (NAR=0.67) and in Figure 6B-H, where g_max_ = 0.75 (NAR=0.67). The maximal conductance of the GABA-A synapses were set to gmax = 1 nS.

A total of 629 excitatory and 120 inhibitory synapses were uniformly distributed across the entire dendritic tree using the following procedure: An excitatory and an inhibitory synapse was placed at the somatic end of each dendritic branch and further excitatory and inhibitory synapses were added at every 10 *μm* and 100 *μm* distances, respectively. An additional set of N_soma_=420 inhibitory synapses were also added to the soma to model the effect of strong perisomatic inhibition.

To account for the possible depolarization caused by the presence of the recording electrode in the experiments, a small (0.02 nA) constant current was injected to the dendritic branch we monitored in the biophysical model during the simulations shown in Fig. 2.

The **cerebellar granule** cell was modelled using a spherical somatic compartment (diameter: 5.8 μm) connected to four 20 *μm* long dendritic branches (diameter: 0.75 μm), an axon hillock (d=1.5 μm, L=2.5 μm) and an axon (d=1 μm, L=200 μm). The passive parameters were *C_m_ = 1* pF/cm^2^, *R_m_* = 5000 Vcm^2^, R_i_ = 100 Vcm. In the simulations with somatic spiking (Figure 7D-E) voltage-activated Na^+^ channels (800 S/cm^2^) and voltage-activated K^+^ channels (20 S/cm^2^ soma and hillock, 15 S/cm^2^ axon) were also inserted.

Each dendritic branch was contacted by one inhibitory (GABA-A and GABA-B) and one excitatory (AMPA and NMDA) synapse, modelled with bi-exponential functions with time constants and maximal conductance of AMPA τι = 0.22 ms, τ_2_ = 2.5 ms, g_max_ = 0.2 nS;NMDA τι = 3 ms, τ_2_ = 40 ms, g_max_ = 0.2 nS;GABA-A τι = 0.39 ms, τ_2_ = 6.8ms, g_max_ = 0.5 nS and GABA-B *τ\* = 30 ms, τ_2_ = 150 ms, g_max_ = 0.2nS (the inhibitory reversal potential was set to −65 mV). The application of Gabazine was modelled by setting the maximal conductance of GABA-A to 0.

### Inputs

We modelled the synaptic input of L23 pyramidal neurons during the presentation of 16 differently oriented moving grating stimuli by the combination of the following three factors: 1) orientation dependent cell assembly dynamics; 2) slow fluctuations in the firing rates; 3) Poisson spike generation. This input structure was chosen to provide a rich stimulus set to engage dendritic non-linearities and match the *in vivo* observed dendritic membrane potential dynamics.

To model presynaptic cell-assembly dynamics the excitatory inputs were divided into 13 orientation-tuned functional assemblies, where neurons within an assembly were correlated with each other and neurons from different assemblies were independent (Figure 4A). Cells from each assembly targeted a particular subtree of the entire dendritic tree (Figure 2I) facilitating the generation of dendritic spikes (Polsky et al., 2004) and implementing synaptic clustering (Wilson et al., 2016). Neurons within an assembly switched randomly and synchronously from a background firing rate (5 Hz) to an elevated activity (20 Hz), where the rate of switching on changed between Ω?η = 0.5 Hz and Ω_on_ = 14 Hz in a sinusoidal manner depending on the orientation of the stimulus. The duration of the active states had an exponential distribution directed by the switching off rate, Q_off_ = 20 Hz, independent of the stimulus orientation. The maximum duration of the active states was set to 150ms. The preferred orientation of each dendritic branch (location of the maximal on-rate) was randomly chosen from a normal distribution with parameters μ = 0^o^ and σ = 33^o^ and then rounded to the nearest multiple of 22.5^o^ (to match the 16 different stimulus orientations).

To generate more smoothly varying inputs and to increase the trial to trial variability characteristic to the experimental data, we also added a slowly decaying fluctuation component to the excitatory firing rate independently for each cell assembly. Specifically, the actual firing rates followed an Ornstein-Uhlenbeck process, decaying towards the state-dependent equilibrium rates (set by the switching process) with a time constant τ = 500 ms and having a standard deviation of 2.5 Hz and 10 Hz in the background and in the elevated state, respectively (Ujfalussy et al., 2015). Finally, spiking of the neurons was modelled as an inhomogeneous Poisson process with the rates defined above.

The inhibitory presynaptic neurons did not show orientation tuning, and were all weakly, but positively correlated with the excitatory neurons (Haider et al., 2013). Their firing rate was proportional to the instantaneous mean of the excitatory firing rates and switched between 20 Hz (when all excitatory assemblies were in the background state) and 30 Hz (all excitatory assemblies being in the elevated state).

Figure 2K-M shows data averaged over 18 s of activity for each of the 16 different orientations. To train the hLN models, we generated 10 different repetitions (with random state transitions, slow fluctuations and spikes) of 48 s long stimulus blocks consisting of 3 s long sections of each of the 16 orientations.

To demonstrate the robustness of our results, we varied either the input firing rates (Figures S5) or the input correlations (Figures S6). In these figures we only modelled presynaptic assembly dynamics, varying the background and the elevated firing rates of the presynaptic excitatory and inhibitory cells (Table 1), but not the orientation selectivity of the inputs or the slow fluctuations in the presynaptic firing rates. The switching on and off rates were Ω_on_ = 1 Hz and Ω_off_ = 10 Hz, respectively.

**Table 1.** The firing rates of the presynaptic neurons in Figure S5.

| label | background (Hz) |  | elevated (Hz) |  | N<sub>soma</sub> |
| --- | --- | --- | --- | --- | --- |
|  | excitatory | inhibitory | excitatory | inhibitory |  |
| 5 | 1 | 5 | 5 | 7 | 420 |
| 10 | 3 | 10 | 10 | 15 | 420 |
| 20 | 5 | 20 | 20 | 30 | 420 |
| 25 | 6 | 55 | 25 | 80 | 100 |
| 40 | 8 | 80 | 40 | 100 | 100 |
| 55 | 10 | 150 | 55 | 300 | 100 |

To analyse the mechanisms underlying input multiplexing, we simulated a simpler scenario where only 160 excitatory and 32 inhibitory synapses distributed on 4 different dendritic branches was stimulated (excitatory, background: 4 Hz, peak: 24 Hz; inhibitory, background: 14 Hz, peak: 60 Hz).

Input for **cerebellar granule cells** *in vivo* during brief air-puff stimulation of the lip area and whiskers (Duguid et al., 2012, 2015) was modelled as an inhomogenous Poisson process with one of the four presynaptic excitatory mossy fibers and inhibitory Golgi cells transiently increasing (mossy fibers: 150 Hz; Golgi cells: 60Hz, duration: 80 ms) and then decreasing (mossy fibers: 6.25 Hz; Golgi cells: 0.75 Hz, duration: 0.4 s) their firing rates compared to a baseline activation (mossy fibers: 6.25 Hz; Golgi cells: 2.5 Hz). The inter stimulus interval was 2 s.

### hierarchical Linear-Nonlinear (hLN) model

To study non-linearity of the input output transformation of neurons we developed the hierarchical linear-nonlinear (hLN) model composed of a cascade of linear and non-linear processing units (Figure 3). Here we describe the details of the model as well as the procedure we used to fit the model to data.

The input spike trains are represented by a vector s(*t*) of length *N* which is the number of presynaptic neurons, indexed by *i*. Each of the *M* dendritic subunit receives input from *M_j_* = {0, N} presynaptic neurons through synapses characterised by their time constants, *τ_ji_*, their propagation delays, *Δ_ji_* and synaptic weights, *w_ji_*. The total synaptic input, *x_j_*(*t*) to dendritic subunit *j* is:

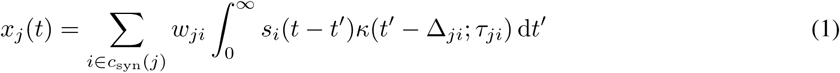

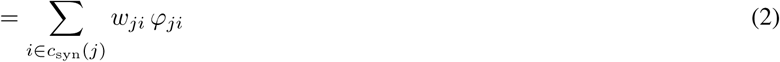

where,φ_ji_ is the total input at a given synapse, *κ*(t; τ_ji_, Δ_ji_) is the synaptic kernel and c_syn_(*j*) denotes the set of indices for the synapses connected to subunit *j*. We used the standard alpha function for synaptic kernels:

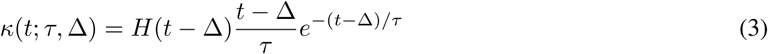

where *H(·)* is the Heaviside step function. We used the combination of two different α-kernels per excitatory synapse for the L2/3 neuron and for both inhibitory and excitatory synapses in the granule cell model. The two kernels were necessary as we found that the functional form of a single alpha synapse was too restrictive to capture linear integration properties of the cells with a mixture of fast and slow synaptic receptors. The two kernels belonged to the same subunit (i.e., sharing the same non-linearity) and captured linear integration at different time scales, and is not considered multiplexing (which requires different non-linearities, see below). The amplitude of the kernels were independent parameters (w^fast^ and w^slow^) but we found that their time constants could be coupled through a simple, linear relationship τ^slow^ = 10.4 + 2.8 τ^fast^ without changing the quality of the fits but decreasing the number of parameters.

When studying input multiplexing (Figure 6), each subunit were allowed to have two different non-linearities each of them associated with one (Figure 6B-H) or two (Figure 6A) α-kernels for each presynaptic spike train.

In Figure 6B-H we allowed only one kernel per non-linearity in order to highlight the differences between the processing channels.

The total input to a given subunit is the sum of the synaptic inputs and the inputs arriving from other connected subunits:

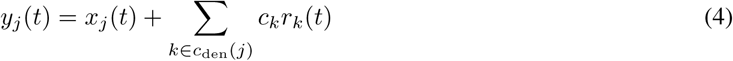

where where c_den_(*j*) denotes the set of indices for the subunits connected to subunit *j*, *c_k_* is the strength of coupling of subunit *k* to its parent and *r_k_*(*t*) is the activation of subunit *k*, which is a (logistic) sigmoid function:

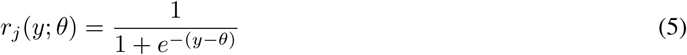

We chose sigmoid for several reasons: First, the sigmoid has been proposed elsewhere as an appropriate dendritic non-linearity (Poirazi et al., 2003a; Polsky et al., 2004). Second, under different parameter settings and input statistics, the sigmoid is flexible enough to capture purely linear, sublinear, and superlinear behavior, as well as combinations thereof. The single free parameter of the sigmoid is its threshold, *θj*, since its slope is set by parameters *wji* and *c_k_* and its output scale is defined by *cj*. Note that the derivatives of the sigmoid function wrt its argument and its parameter both have a particularly simple form:

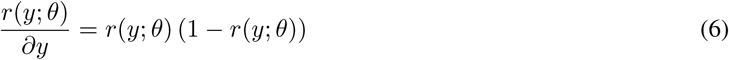

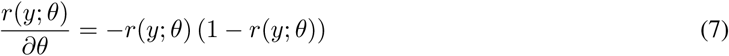

In some simulations the sigmoid non-linearity was omitted from the somatic compartment (e.g., Figure 4F, left) leading to linear integration.

Finally, in the simulations shown in Fig. S4 the model was extended to incorporate postsynaptic spiking (Mensi et al., 2012) which leads to a hierarchical Generalised Linear Model (hGLM). Spiking of the postsynaptic neuron triggered adaptation currents which we modeled with an extra additive term in the membrane potential of the somatic subunit. We used a set of N_*ψ*_ = 10 basis functions of raised cosine ??bumps??, each convolved with the *output* spike train leading to adaptation kernels *ψ_i_(t)*:

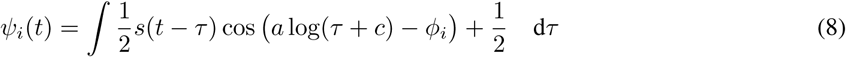

for τ such that *a* log(τ + *c*) * *{ϕ_i_ — π, ϕ_i_* + π} and 0 elsewhere, and *a* = 3.75, c = 0.01 and *ϕ* set uniformly in the interval (3,22) (Pillow et al., 2008). The total effect of the postsynaptic spikes on the somatic membrane potential was the sum of the adaptation kernels weighted by the coefficients *ϒ_i_*.

In summary, the response of the hLN model to synaptic inputs is given by

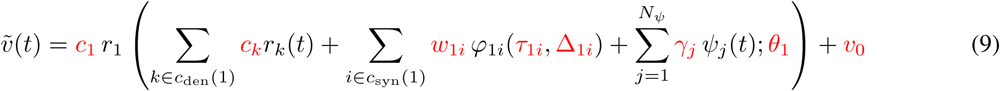

where the response 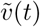 corresponds to the subthreshold somatic membrane potential of the cell, and subunit *k* = 1 refers to the soma, which is the root of the hierarchy. Importantly, both the somatic response and its derivative wrt. the parameters can be evaluated in a single sweep starting from the leaves (terminal subunits) and ending at the root subunit.

### Model fitting procedure

The goal of the model fitting was to match the biophysical model’s somatic membrane potential, v(*t*) with the response of the hLN model, 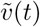, to the same input spike trains.

We assume in this work that the hLN architecture, defined by the lists c_syn_ and c_den_, is given in advance, that is, instead of learning the structure of the model we can chose from a couple of candidates based on their ability to predict test data. During fitting we minimise the fitting error 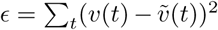, the squared deviation between the subthreshold component of the training signal and the hLN model’s response using gradient descent. To avoid local minima we first train simple models and use them to initialise the parameters of more complex models. By using this procedure, simpler models also provide an upper bound on the training error for the more complex models.

Specifically, we initially coupled the parameters of all synaptic kernels such that all synapses shared a common amplitude, time constant and delay. Moreover, we always started with a single subunit-model, which substantially reduced the number of parameters to be fitted and helped avoiding shallow local optima. Next, we initialized models with hierarchical subunit structure and one synapse per subunit by pre-tuning the non-linearities of the subunits to approximate linear integration. Specifically, we rescaled the inputs to the subunits by changing the parameters *w_ji_* and *c_k_* such that *n*-SD of the total input overlapped with the central, approximately linear part of the sigmoid non-linearity. We repeated this scaling with various values of *n* and chose the one which resulted in the lowest training error after optimization.

Finally we decoupled the parameters of the synaptic kernels from each other: synapses within each dendritic branch were divided into three groups based on the location of the synapse relative to the tip of the branch (proximal, middle, distal) and only synapses within each group had identical parameters, whereas the time constant and the amplitude of the synaptic kernels was allowed to vary between groups (we used a single synaptic delay for each subunit even in the unconstrained case). Note that the number of synapse-groups is determined by the morphology of the cell and is independent of the number of subunits.

In order to prevent overfitting, we used a log-normal prior for the individual response amplitudes and the synaptic time constants. The mean parameter of the prior was set by the parameter values found by the coupled optimization, and the variance parameter was set to match to the variance of the somatic PSPs recorded in response to individual synaptic stimulations in the biophysical model. Note, that in a hLN model the response amplitude of synapse i targeting subunit *j* is 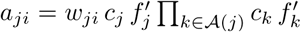, where 𝒜 (*j*) denotes the ancestors of subunit *j*, i.e., subunits towards the root. Importantly, our prior on *a_ji_* imposes constraints not only on the synaptic wight but also on the resting slope of the subunit non-linearity as well as on the subunit couplings.

Parameters of the hLN model were fitted and evaluated on 10 separate segments of 40 s long training and test data. We quantified the accuracy of the models by the variance explained in the subthreshold signal, which is the fitting error normalised by the variance of the signal: ε = 1 - ε/Var[*𝜐*(*t*)]. Parameters of the spiking response were fitted using a standard maximum likelihood approach after fitting the parameters subthreshold response (Mensi et al., 2012) and the fits were evaluated by the spike prediction accuracy in 3 ms time window.

### Data analysis

Our current analysis of *in vivo* dendritic activity is based on a single recording from Smith et al. (2013) that showed clear orientation tuning in its response but not anaesthesia-related artifacts (i.e., strong up and down states in the absence of the stimulation). Our analysis focused on the 3 s long stimulus periods during each of the 6 repetition of the 16 different stimulus orientations. The plateau probability was calculated as the fraction of time the dendritic membrane potential was above −35 mV. Details of the experimental recording protocol and stimulus presentation can be found in Smith et al. (2013).

The expected dendritic response of the biophysical model (Figures S5D-F, Figures S6D-F) was calculated by first summing the local dendritic PSPs recorded during individual synaptic stimuli and next linearly scaling the sum to best match the measured response during *in vivo*-like stimulation. The number of dendritic Na^+^-spikes (Figures S5I and Figures S6I) was determined by counting the number of upward zero crossings of the local dendritic membrane potential in the biophysical model.

## Acknowledgements

We are grateful to Judit K Makara for helpful discussions and support; Spencer L Smith for providing the *in vivo* data; DJ Strouse for discussions and his contribution in designing the first version of the hLN model; This work was supported by an MTA postdoctoral fellowship (B.B.U.), and the Wellcome Trust (T.B., M.L.).

## Author Contributions

B.B.U., T.B. and M.L. designed the study. B.B.U. performed the simulations and analysed the data. M.L. and T.B. supervised the project and all authors contributed to writing the manuscript.

## Supplemental Figures

**Figure S1.**
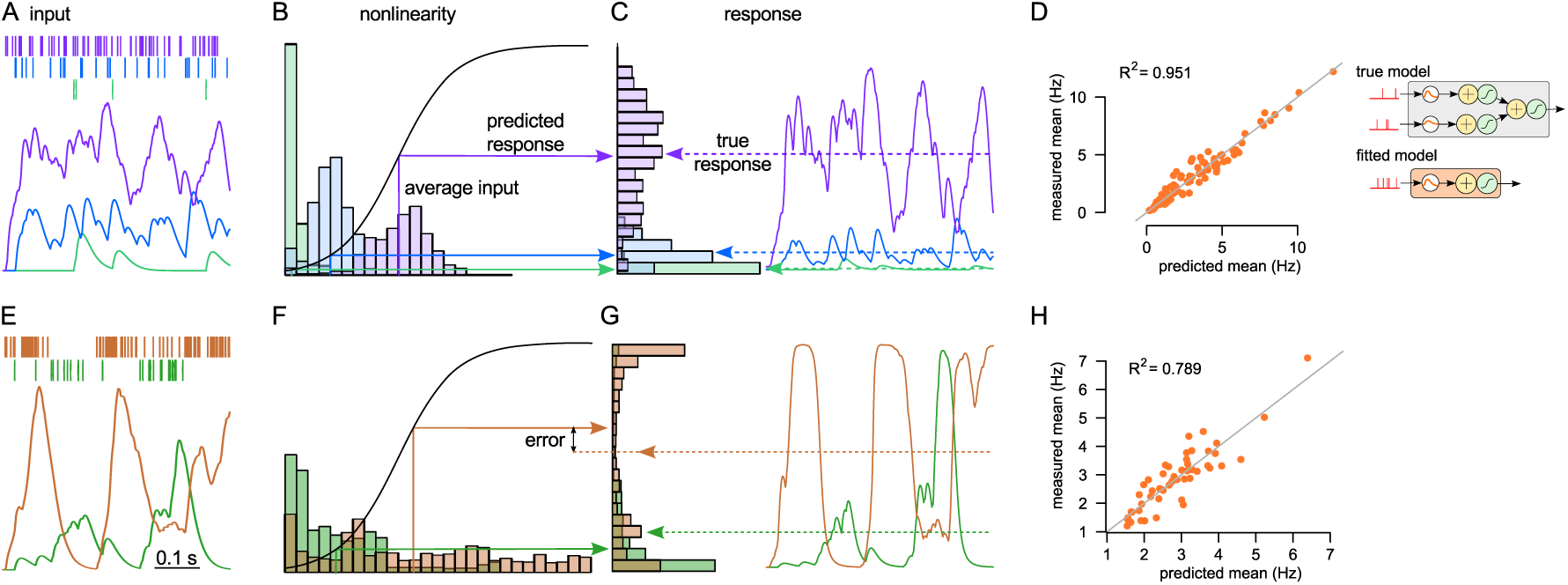
Constant firing rate inputs can mask true dendritic nonlinearities. (A-C) The average response of an LN subunit is accurately predicted by the average input when the inputs have constant firing rates (homogeneous Poisson processes). Input spike trains with different firing rates (A, top, colored rasters) are filtered through a synaptic kernel (A, bottom, filtered traces) and transformed by a sigmoid non-linearity (B, black curve) to produce the response (C, colored traces). Colored histograms show the distribution of inputs (B) and responses (C). Colored arrows show the response predicted by the average input (B) and the actual average response (C). As each input uses only a modest range of the non-linearity, the average input accurately predicts the average response. (D) To illustrate the accuracy of firing rate prediction for constant input firing rates, we stimulated a 2-layer hLN model (inset, top) with a wide range of input firing rates (5-500 Hz) held constant over each 1-s trial, and measured its average output firing rate over the trial. We fitted a one-layer model without local nonlinearities (but with a global non-linearity, inset, bottom). We found that this simple model, implying linear dendritic processing, predicted the response of the true model, including strong local nonlinearities, with high accuracy (R^2^ = 0.95). (E-G) Same as A-C but predicting the response of an LN unit to fluctuating stimuli. When the inputs show strong fluctuations (E), the average input (F) is not predictive of the average output (G), resulting in a substantial prediction error (F-G, arrows). (H) Same as D but for fluctuating inputs. The one-layer model is less accurate in predicting the firing rate of the same 2-layer hLN model as in D (R^2^ = 0.79). Fluctuating input was generated by randomly resampling the input rate from the 5-500 Hz interval every 100 ms. In the simulations shown in panels D and H a constant 150 Hz background input was also added to model noisy input processing or unobserved stimuli.

**Figure S2.**
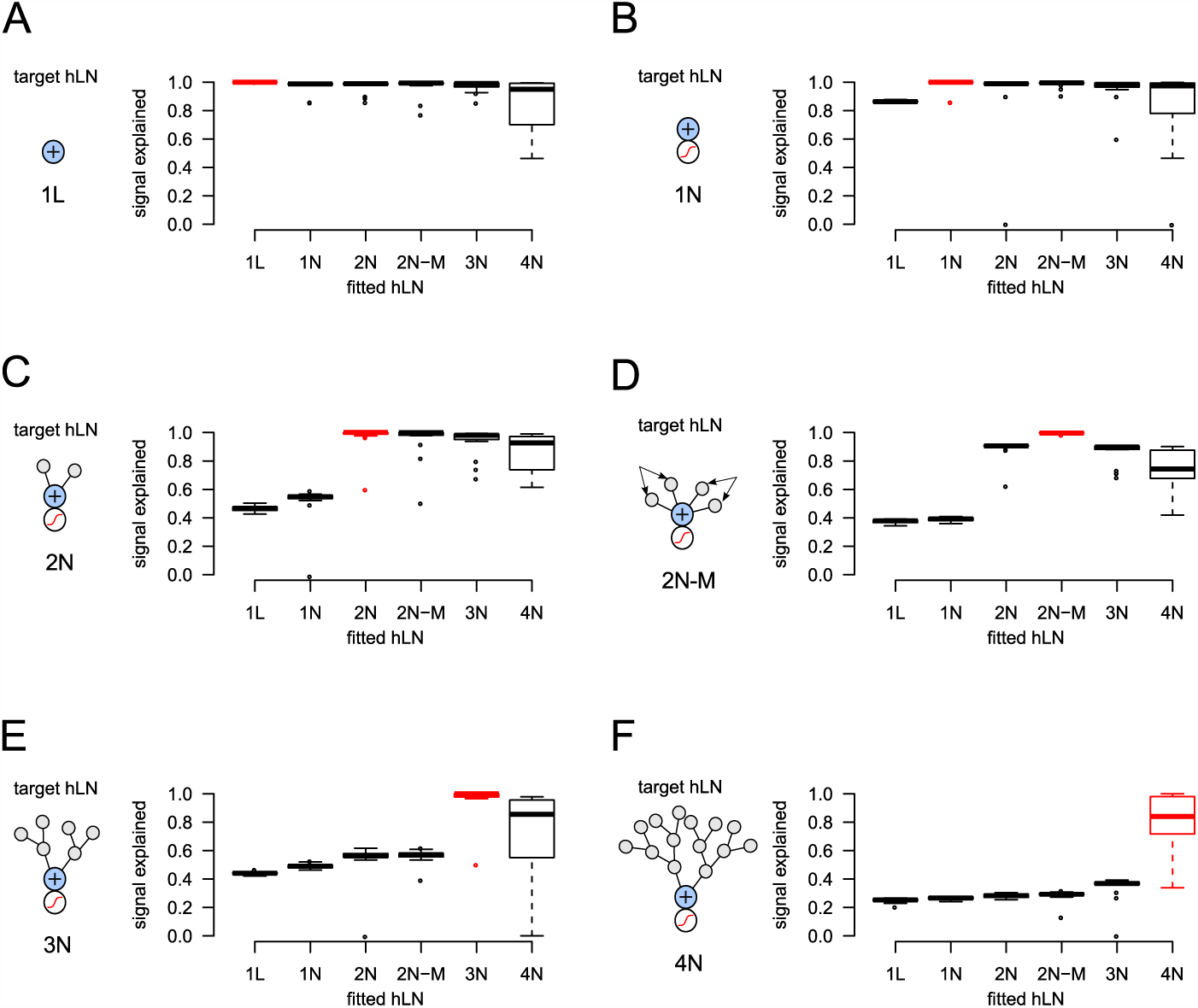
Validation of the model fitting. We generated synthetic data by simulating the response of target hLN models with increasingly more complex architecture (the schamatics in panels A-F; the name of the model indicates the number of layers in the hierarchy (1-4), the presence or absence of an output non-linearity (L or N) and multiplexing dendrites, M in panel D) to random input spike trains. We fitted this synthetic dataset (composed of the input - response pairs of the hLN models) using hLN models with various architectures (x-axis, the same set of models as used to generate data) starting from random initial parameters. The quality of the fit was evaluated by the signal explained on held out test data. We found that in most of the cases the structure of the original model could be correctly recovered by taking the simplest amongst models achieving maximal performance, indicated by the red color. Boxplot shows median, quartiles and the range of 20 independent simulations.

**Figure S3.**
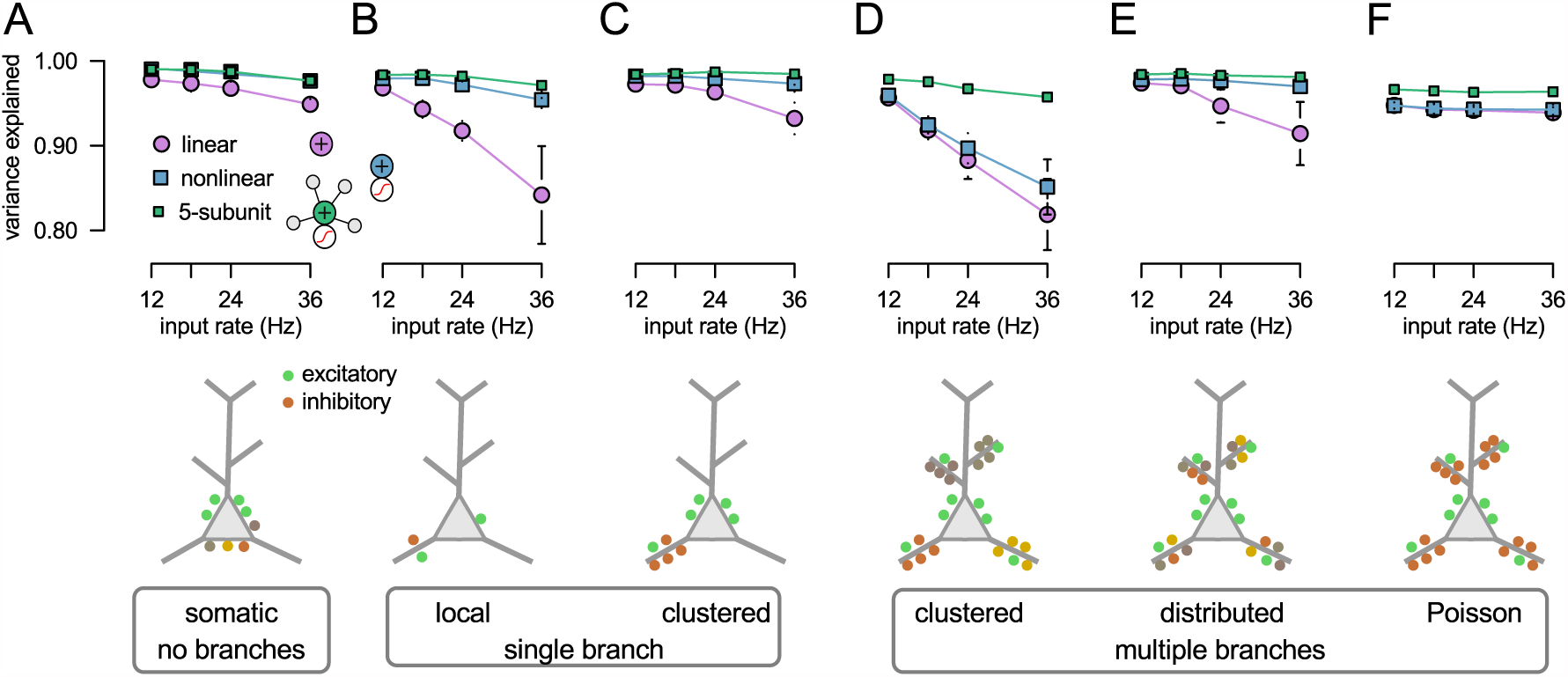
Validation of the model’s assumptions. To rigorously validate the ability of the hLN model class to correctly capture the integration of spatially distributed inputs, despite its drastic discretization of the cell’s morphology into a small number of independent subunits, we conducted a series of simulation experiments varying the spatio-temporal complexity of the inputs. Specifically, we varied the location of the inputs as follows (see schematics below A-F): (A) all excitatory and inhibitory synapses were placed at the soma; (B) a single presynaptic ensemble targeted the same location within a single dendritic branch; (C) similar to B, but synapses were distributed along the whole length of a single dendritic branch; (D) four presynaptic ensembles targeted 4 dendritic branches in a clustered way: neurons from the same ensemble were distributed along the whole length of a single dendritic branch (as in Fig. 6B); (E) similar to D, but inputs were randomly assigned to branches; (F) four branches were targeted, but the inputs were uncorrelated (Poisson spike trains with constant firing rate). Synaptic input patterns were similar to the inputs used in Fig. 6B-H, except that in each conditions we varied the peak excitatory firing rate between 12 and 36 Hz and we used only a single presynaptic ensemble in panel B and uncorrelated inputs in panel F. The response of a passive biophysical model expressing only NMDA non-linearities was fitted by three different hLN architectures (insets in A and B): a point neuron with linear output subunit (linear); a point neuron with nonlinear output subunit (non-linear); and a neuron with 4 dendritic and 1 nonlinear output subunit (5-subunit). Note, that for panels A-C some subunits of the 5-subunit model does not receive inputs. We found that the model in which the subunits corresponded to individual dendritic branches always achieved excellent performance (> 90% variance explained). Specifically, the assumption that dendritic branches correspond to discrete integrative compartments (subunits) is justified by the similar fraction of variance is explained when inputs are distributed along the whole dendritic branch (C) versus when they target a single location within the dendritic branch (B) or when dendritic non-linearities are bypassed by placing all synapses directly to the soma (A). Moreover, the assumption of unidirectional coupling between the subunits is supported by the similar fraction of variance explained by the 5-subunit model when inputs are targeting a single (C) versus multiple dendritic branches (D-F).

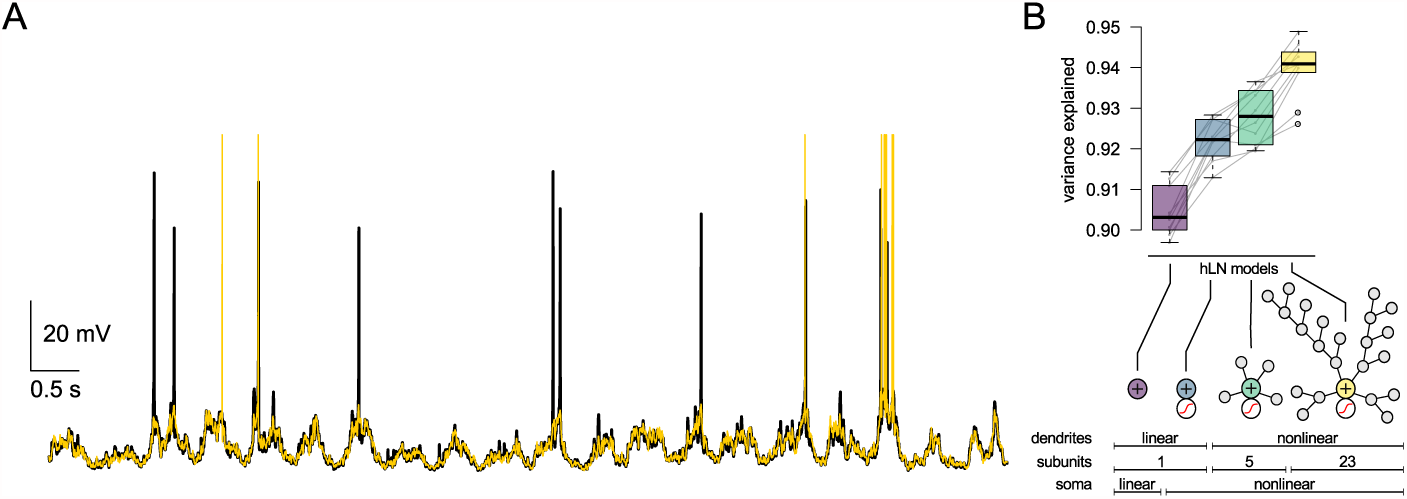
Model with somatic spiking. (A) Somatic membrane potential response of the biophysical model L23 neuron to the same input as in Figure 4 with somatic spiking conductances (black) and the sub- and suprathreshold activity predicted by a hierarchical model with 23 subunits. (B) Subthreshold variance explained by different hierarchical models when predicting the activity of a biophysical model with somatic spiking.

**Figure S5.**
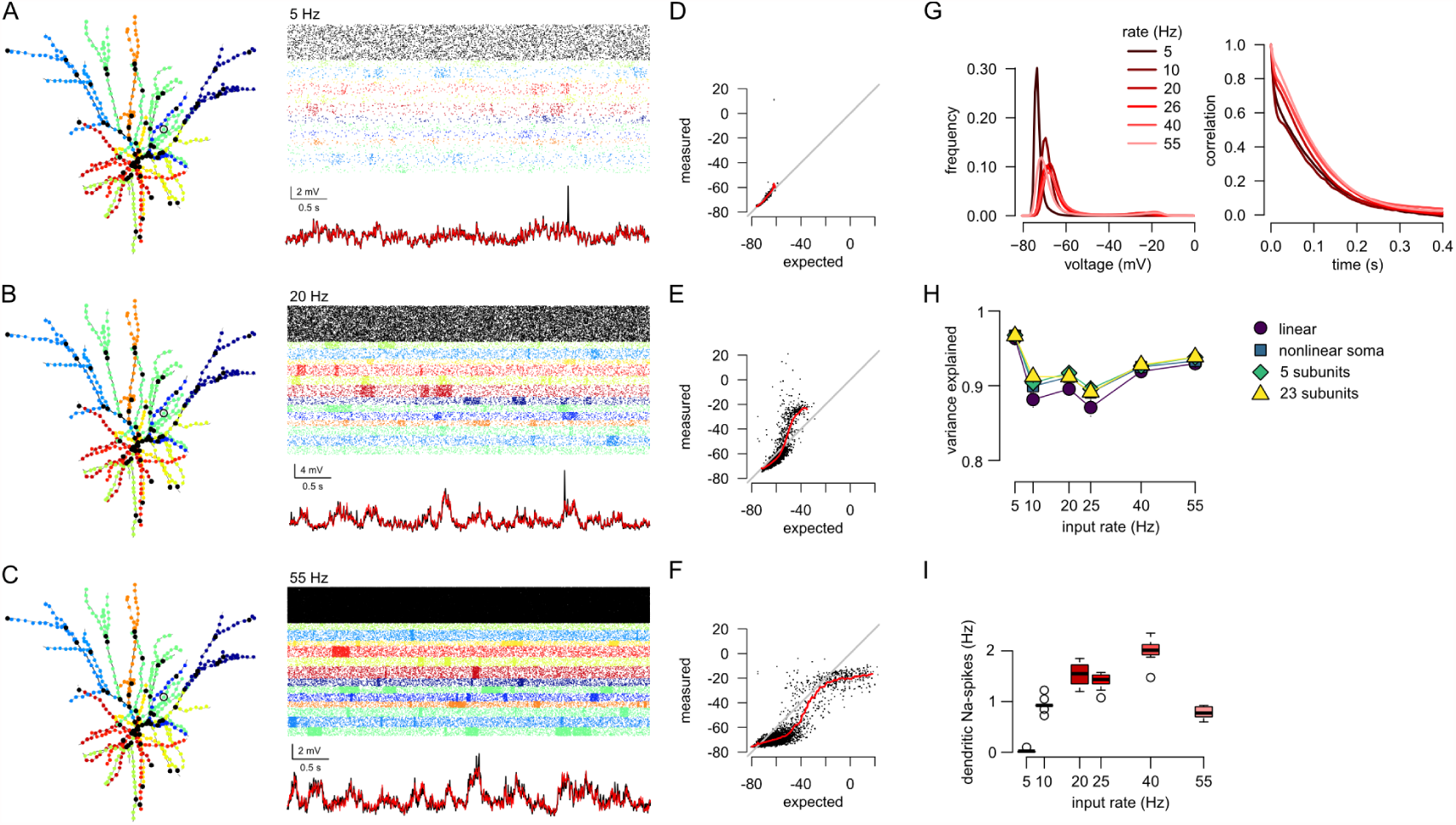
Dependence of dendritic input-output transformation on input clustering. (A-C) Synaptic clusters (left, color coded) and synaptic inputs (top, color code as in Figure 4A) and somatic output (bottom, black) with excitatory input peak firing rates increasing from 5 to 55 Hz (Table 1). Red line indicates the response predicted by a hLN model with 23 subunits. (D-F) Measured dendritic membrane potential as a function of the expected response. Black dots: individual synapses, red line: running average trend line. (G) Histogram (left) and auto-correlation (right) of the dendritic membrane potential at different input rates (color). In the biologically relevant input range the histogram is bimodal and the autocorrelation has a slow decay (cf. Figure 2F-G). (H) Variance explained by hLN models with increasing complexity (same as in Figure 4C) as a function of the input firing rate. (For reference, simulations in Figures 2, 4 and Figure S6 used ~20 Hz input rates.) The performance of linear models are high even when the dendrites show substantial nonlinearity (e.g., panel F). (I) Frequency of dendritic spikes in a representative dendritic branch of the biophysical model as a function of the input firing rate.

**Figure S6.**
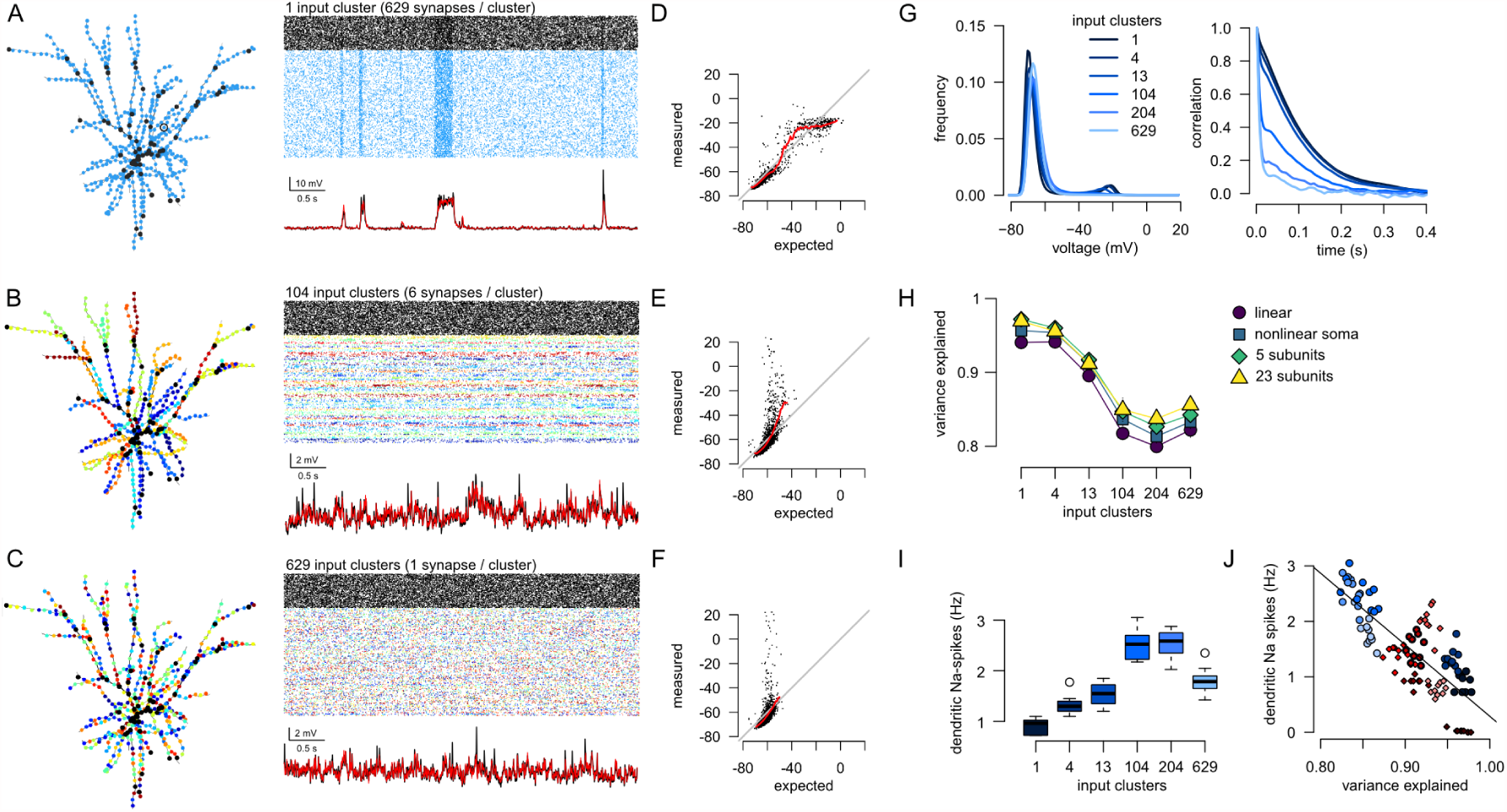
Dependence of dendritic input-output transformation on input clustering. (A-C) Synaptic clusters (left, color coded), and inputs (top) together with the somatic output (bottom) with the number of synaptic clusters increasing from 1 (all synapses belong to the same cluster) to 629 (all synapses are independent). Red line indicates the response predicted by a hLN model with 23 subunits. (D-F) Measured dendritic membrane potential as a function of the expected response. Black dots: individual synapses, red line: running average trend line. (G) Histogram (left) and auto-correlation (right) of the dendritic membrane potential at levels of synaptic clustering (color). In the biologically relevant input range the histogram is bimodal and the autocorrelation has a slow decay (cf. Figure 2F-G). (H) Variance explained by hLN models with increasing complexity (same as in Figure 4C) as a function of synaptic clustering. (For reference, simulations in Figures 2, 4 and Figure S5 contained 13 clusters.) The performance of linear models are high even when the dendrites show substantial nonlinearity (panel D). (I) Frequency of dendritic Na^+^-spikes in a representative dendritic branch of the biophysical model as a function of synaptic clustering. (J) Frequency of dendritic Na^+^-spikes in a representative dendritic branch of the biophysical model as a function of the variance explained by the 23-subunit hLN model for different values of synaptic clustering (shades of blue, as in panel I) and input firing rate (shades of red, as in Fig. S5I). The strong negative correlation indicates that most of the unexplained variance arises from dendritically evoked Na^+^-spikes appearing as spikelets of variable amplitudes in the soma (see also the voltage traces in panels A-C and Figure S5A-C).

